# Hippocampal transformations occur along dimensions of memory interference

**DOI:** 10.1101/2025.10.13.682242

**Authors:** Soroush Mirjalili, Guo Wanjia, Dominik Grätz, Eric Z. Wang, Ulrich Mayr, Brice A. Kuhl

**Affiliations:** Department of Psychology, University of Oregon, 1451 Onyx St., Eugene, OR 97403, USA; Institute of Neuroscience, University of Oregon, 1585 E 13th Ave., Eugene, OR 97403, USA; Department of Psychology, Princeton University, South Drive, Princeton, NJ 08540, USA; Department of Psychology, University of Southern California, Seeley G. Mudd Building, 3620 McClintock Ave, Los Angeles, CA 90089, USA

## Abstract

The role of the hippocampus in resolving memory interference has been greatly elucidated by considering the relationship between the similarity of visual stimuli (input) and corresponding similarity of hippocampal activity patterns (output). However, these input-output functions can take surprisingly different forms. Here, we reconcile seemingly conflicting findings by considering the possibility that the hippocampus prioritizes different dimensions of visual similarity across different stages of learning. First, we generated a set of natural scene images from two visual categories and rigorously characterized visual similarity using a wide array of methods: neural networks, artificial intelligence models, and human perceptual and memory decisions. We then identified two orthogonal dimensions of visual similarity that each predicted memory interference, but that did so at distinct stages of learning. Using high-resolution fMRI, we then tested for dimension-specific input-output functions within the hippocampus. Within CA3 and dentate gyrus (CA3/DG), we show that dimensions of visual similarity were inverted (repulsion) at stages of learning when they contributed to memory interference. In contrast, CA3/DG preserved similarity when dimensions did not contribute to interference. These findings reveal that hippocampal representations of visually-similar stimuli are highly dynamic and critically depend on the dimensions of similarity that currently contribute to memory interference.

## INTRODUCTION

A critical facet of the hippocampus’ role in episodic memory is its capacity to minimize interference^1–6^. Because memory interference is driven by similarity between stimuli^7–11^, theories of interference resolution in the hippocampus have focused on how stimulus similarity (input similarity) translates to the similarity of corresponding hippocampal representations (output similarity). While a general principle of interference resolution is that output similarity should be lower than input similarity^1,5^, this principle can be satisfied in qualitatively distinct ways. Indeed, in some contexts, increases in stimulus similarity yield positive, but sub-linear increases in hippocampal similarity^1,5,12^. In other contexts, increases in stimulus similarity yield no change in hippocampal similarity^13,14^. In yet other contexts, higher stimulus similarity has been associated with *lower* hippocampal similarity^15–17^. A major challenge for theories of the hippocampus is to explain this variability in the shape of input output functions.

One reason why relationships between stimulus similarity and hippocampal similarity may be so variable is because similarity, itself, can be defined using multiple metrics^18–23^ that potentially capture multiple underlying *dimensions* of similarity^18–21,23–25^. This raises the obvious question: which dimensions ‘matter’ to the hippocampus? If a primary function of the hippocampus is to minimize interference, then a simple, but potentially clarifying perspective is that the hippocampus will target dimensions of similarity that drive interference. That said, an underappreciated point is that interference may be a *moving target*: the specific dimensions of similarity that drive interference may shift with learning^26^. While speculative, this may explain why relationships between stimulus similarity and hippocampal similarity sometimes change with experience^16,17,27,28^. However, formally testing this idea requires (a) identifying distinct dimensions of stimulus similarity, (b) characterizing how these dimensions influence memory interference across stages of learning, and (c) testing whether dimension-specific influences on behavior explain the shape of input-output functions in the hippocampus.

Here, we first developed a set of natural scene images from two visual categories and obtained 10 different metrics of visual similarity. These metrics of visual similarity were generated from new human behavioral data and various computational methods. By applying Principal Component Analysis (PCA) to the similarity metrics, we identified two orthogonal dimensions of similarity that strongly predicted memory interference errors. Strikingly, we found that the relative influence of these dimensions on behavior sharply changed with experience (learning). Whereas one dimension drove interference during early stages of learning, the other drove interference during later stages of learning. Finally, using high-resolution fMRI, we show that dimension-specific input-output functions in CA3 and dentate gyrus (CA3/DG) assumed qualitatively different shapes across stages of learning. Specifically, CA3/DG *inverted* each dimension of visual similarity (negative input-output functions) when it contributed to memory interference but preserved each dimension (positive functions) when it was not actively contributing to interference. Thus, entirely opposite coding schemes of similarity dimensions in the hippocampus can be explained by the extent to which these dimensions were behaviorally relevant. These findings provide important insight into how the hippocampus resolves memory interference, demonstrating highly dynamic and adaptive processes that dramatically increase the representational distance between memories that are most at risk for interference.

## RESULTS

### Dimensions of visual stimulus similarity

We first generated a set of 48 visual stimuli (natural scenes) drawn from two categories: beaches and gazebos (**Fig. 1a**). We then selected 10 different similarity metrics that yielded (for each metric) a matrix (**Fig. 1b**) describing the similarity of each pair of scenes within each category (276 beach pairs + 276 gazebo pairs = 552 pairs in total). Two of these metrics were based on human behavioral ratings that we collected in separate experiments. In one experiment (*behavioral memory experiment*), we measured associative memory confusability (**Fig. 1c**, left; also see **Supplementary Fig. 1**). In the other experiment (*perceptual similarity experiment*), we measured subjective judgments of perceptual similarity (**Fig. 1c**, right). The other 8 metrics were based on neural network models [three different layers of VGG-16, AlexNet, and Contrastive Language-Image Pretraining (CLIP)], Natural Language Processing (NLP) applied to AI-generated text descriptions of images (**Supplementary Fig. 2**), a measure of structural image features [Structural Similarity Index (SSIM)], and a model of global image features (GIST). (See *Methods* for details of each measure). Rather than identifying a single measure or model that ‘best’ describes stimulus similarity, our goal was to extract *dimensions* of similarity from the *set* of measures/models.

**Fig. 1.**
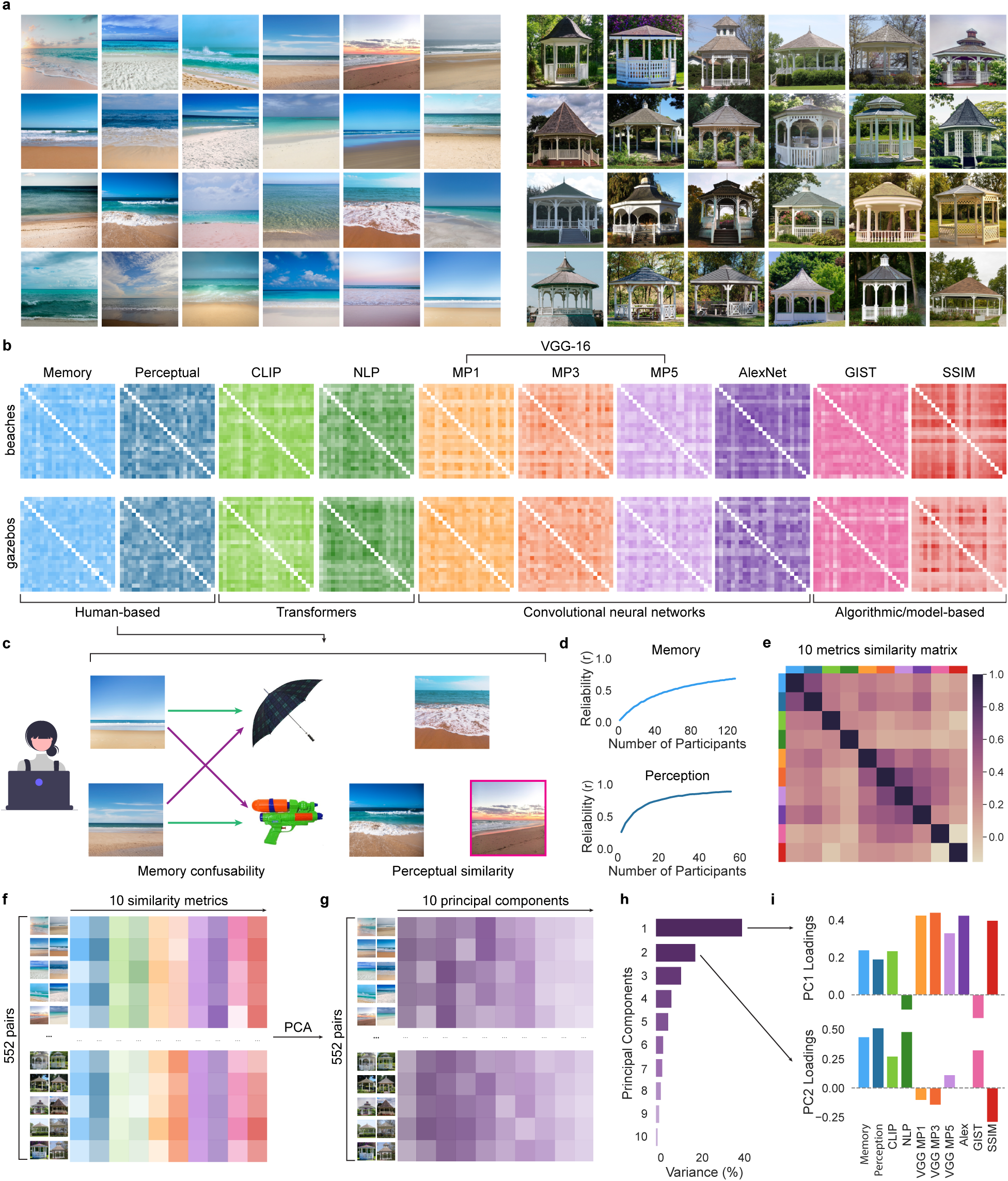
Identifying orthogonal dimensions of visual similarity. **a**, The stimulus set comprised 24 beaches and 24 gazebos (all shown here). **b**, 10 metrics of similarity were selected (see Methods). For each metric, a similarity matrix was generated for each visual category, with each cell representing the similarity between a pair of scenes. **c**, Left, the behavioral memory experiment involved a scene–object associative learning task that provided a measure of memory confusability. Green arrows reflect correct associations; purple arrows represent errors (interference) that were used to generate a confusion matrix. Right, the perceptual similarity experiment used an odd-one-out task, in which similarity between two scenes was defined by the probability that a third scene was chosen as ‘odd’. **d**, Split-half reliability (collapsed across beaches and gazebos) as a function of number of participants included in the behavioral memory (top) and perceptual similarity (bottom) experiments. **e**, Matrix reflecting pairwise similarity (Pearson correlations) of the 10 similarity matrices. Higher values indicate greater alignment of the similarity matrices across metrics. **f**, All similarity data was combined into a single matrix with 552 row (each within-category pair of scenes) and 10 columns (each similarity metric). Each cell represents the similarity between a specific pair of scenes according to a specific metric. **g**, PCA was applied to the matrix shown in **f** to obtain orthogonal dimensions of similarity. Each cell represents the principal component (PC) score for a specific pair of scenes according to a specific PC (dimension). **h**, The variance explained by each PC. **i**, Contribution (loading) for each similarity metric for PC1 and PC2.

Given that the behavioral memory experiment and the perceptual similarity experiment were based on human responses, it was important to establish that resulting similarity metrics were reliable. To test this, we computed split-half reliability. Specifically, we divided the pool of participants in half, calculated similarity matrices within each half of the participants, and then correlated these similarity matrices. For both experiments, the mean split-half reliability (r) exceeded 0.69, with reliability asymptotically increasing as a function of the total number of participants included (**Fig. 1d**).

While the 10 similarity matrices were generally positively correlated with each other, Pearson correlations ranged from −0.154 to 0.665 (**Fig. 1e**). To identify different dimensions of similarity, each of the 10 similarity matrices was vectorized and combined in a new matrix that contained the pairwise similarity for all 552 within-category pairs across all 10 similarity metrics (**Fig. 1f**). We then applied a principal component analysis (PCA) to this new matrix to identify orthogonal components (dimensions) of similarity across the 10 metrics (**Fig. 1g**). The first 2 principal components (PC1 and PC2) explained over 60% of the variance in the similarity matrix (41.3% and 19.1%, respectively; **Fig. 1h**). For PC1, the metrics with the highest positive loadings were VGG max pooling layers 1 and 3, SSIM, and AlexNet. For PC2, the metrics with the highest positive loadings were memory confusability (behavioral memory experiment), perceptual judgments (perceptual similarity experiment), NLP of text descriptions, and GIST (**Fig. 1i**). These loadings suggest that PC1 captured low- to mid-level perceptual similarity, whereas PC2 captured semantic similarity (see *Methods* for additional consideration of PC interpretations and see **Supplementary Fig. 3** for pairs of images with high vs. low PC scores). The loadings for all 10 PCs are in **Supplementary Table 1**.

### Establishing behaviorally relevant dimensions of similarity

We next sought to establish the dimensions of visual similarity that were related to behavioral expressions of memory interference. We addressed this by analyzing behavioral data from the *fMRI experiment*, which was entirely independent from the data reported above. The fMRI experiment was modeled after the behavioral memory experiment (**Fig. 1c**), using the same stimuli (24 beaches, 24 gazebos) and associative learning framework. The experiment included three phases (**Fig. 2a**): scene-object associative learning conducted prior to fMRI scanning (training phase), scene exposure during scanning (exposure phase), and an associative memory test conducted after scanning (post-test).

**Fig. 2.**
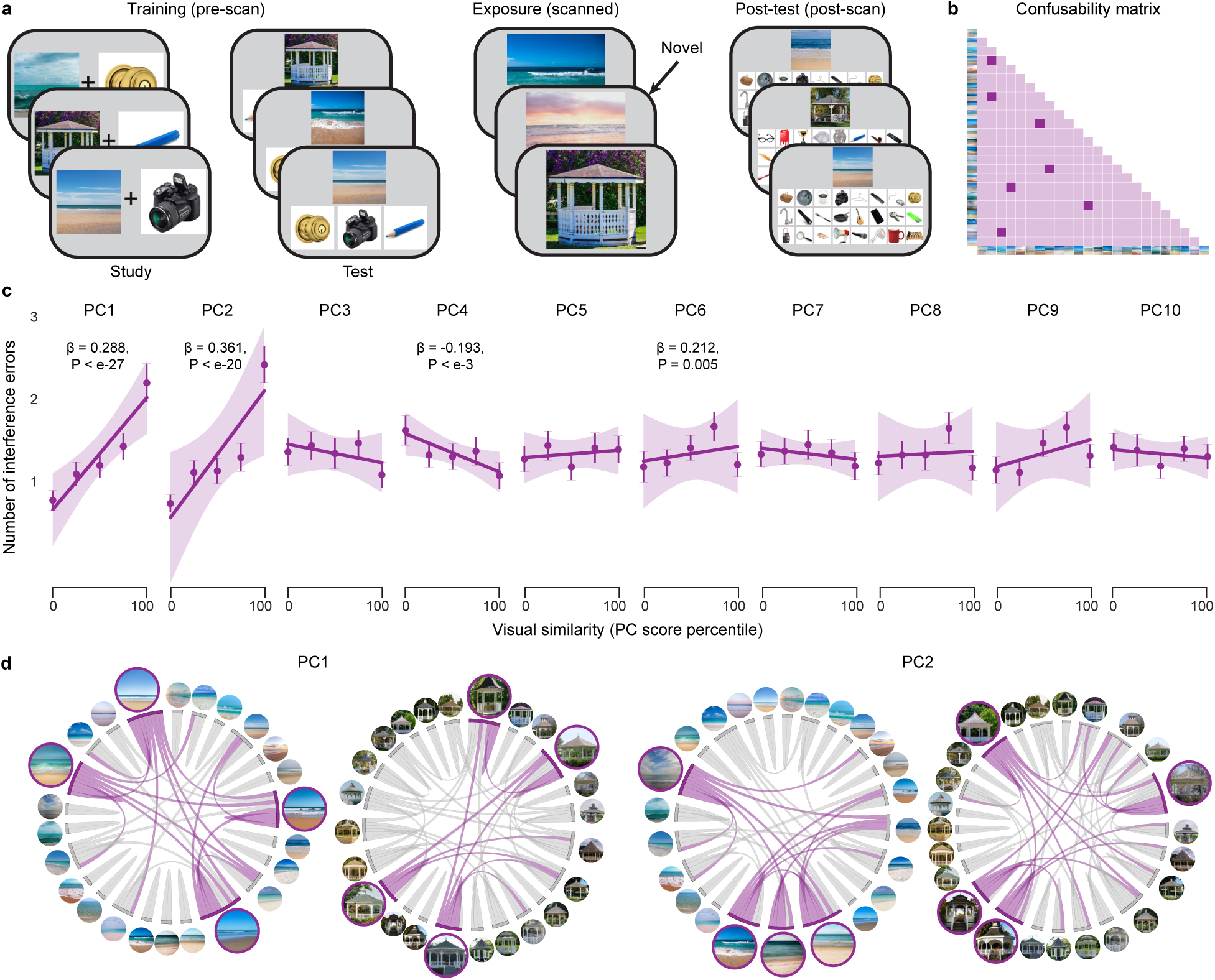
Establishing behaviorally relevant dimensions of similarity. **a**, Overview of the fMRI experiment: participants completed scene–object associative learning before scanning (training phase, left), a novelty detection task during scanning (exposure phase, middle), and an associative memory test after scanning (post-test; right). **b**, Post-test errors yielded a confusability matrix for each participant, indicating which pairs of scenes were confused. **c**, Relationship between visual similarity (PC scores derived as shown in Fig. 1) and pairwise confusability in the post-test of the fMRI experiment. Statistical analyses (linear mixed-effects models) treated similarity as a continuous variable. Lines of best fit (solid lines) and S.E.M.s (shaded areas) are shown. Circle markers show the mean number of errors for 5 levels (bins) of similarity. Error bars indicate S.E.M. These binned data are purely for visualization purposes. For similarity dimensions (PC scores) that significantly predicted the number of interference errors, the corresponding beta estimates and p-values from the mixed-effects models are shown. **d**, Circle graphs indicating the similarity between scenes from each category based on PC1 score (left) and PC2 scores (right). Edge thickness reflects the similarity between a given pair of scenes; only the most strongly connected pairs—those with similarity above the cut-off threshold (0.65–0.72 across the four graphs)—are shown for clarity. For each graph, the four scenes with the highest degree centrality (aggregate connection strength) are shown with larger node sizes and a purple border.

The training phase consisted of alternating study-test rounds (3 total). During study rounds, each scene was presented with a unique object. During test rounds, scenes were presented along with three objects and participants selected the associated object. Of the three objects, one was correct (target), one was associated with a scene from the same category as the current scene (competitor) and one was associated with a scene from the other category (non-competitor). Importantly, for each participant, one visual category was designated the ‘high training’ category, and the other was ‘low training’. High training associations were presented more often, for longer durations, and with feedback during test rounds. During the exposure phase, participants were repeatedly shown all 48 scenes from the training phase, along with a handful of novel scenes. Participants were instructed to identify the novel scenes. In the post-test, each scene was presented along with all 24 objects that had been associated with that category (1 target, 23 competitors). Participants attempted to select the target.

The post-test data were used to identify dimensions of visual similarity that produced interference. In particular, the data were used to generate a pairwise memory confusability matrix for each visual category (**Fig. 2b**). To identify similarity dimensions that predicted memory interference errors, we ran mixed-effects models (a separate model for each PC) for which the dependent measure was the pairwise value from the post-test confusability matrix and the predictor included PC scores for one of the 10 dimensions of similarity (**Fig. 1g**). Significant effects of PC score (at Bonferroni corrected α = 0.005) were observed for PC1 (β = 0.288, t_(28699)_ = 10.933, P < 0.001), PC2 (β = 0.361, t_(28699)_ = 9.470, P < 0.001), PC4 (β = −0.193, t_(28699)_ = −3.478, P < 0.001), and PC6 (β = 0.212, t_(28699)_ = 2.823, P = 0.005) (**Fig. 2c**). As a robustness test, we also ran an alternate version of the analysis in which we computed subject-level correlations (Pearson correlation) between confusability matrices and PC-derived similarity matrices. Resulting correlation coefficients were z-transformed and compared against zero via one-sample, two-tailed t-tests. Significant effects of PC score (at Bonferroni corrected α = 0.005) were observed for PC1 (t_(51)_ = 10.251, P < 0.001, d = 1.450), PC2 (t_(51)_ = 10.672, P < 0.001, d = 1.509), and PC4 (t_(51)_ = −3.422, P = 0.001, d = −0.484).

Given that PC1 and PC2 explained over 60% of the variance in the combined similarity matrix (**Fig. 1h**) and they each strongly predicted behavioral interference errors, subsequent analyses focused on these two components. For visualization, **Fig. 2d** shows the similarity structure of all scenes within each category, separately for PC1 and PC2. In this visualization, connections between pairs of images are equivalent to edges in a graph and individual images are equivalent to nodes. Pairwise similarity between images can therefore be described in terms of edge strength whereas an individual stimulus’ aggregate similarity to all other images can be described in terms of degree centrality.

### Experience-dependent changes in behavior

The training manipulation in the fMRI experiment (high vs. low training) was used to test for experience-dependent changes in behavior and the hippocampus. To confirm that the training manipulation was successful, we used mixed-effects models to assess behavioral performance during the training phase (prior to scanning), the exposure phase (during scanning), and the post-test (after scanning). During the training phase, there was a main effect of training level (high vs. low; β = 0.347, t_(307)_ = 12.313, P < 0.001; **Fig. 3a**, left) and a main effect of test round on memory accuracy (β = 0.088, t_(307)_ = 11.460, P < 0.001). These main effects indicate, respectively, that associative memory was (i) better with high vs. low training and (ii) improved across test rounds. When participants did make errors during the training phase, these tended to be ‘interference errors’: choosing an object that was associated with the same scene category as the target, as opposed to the opposite category (β = 0.231, t_(615)_ = 15.091, P < 0.001; **Fig. 3a**, right). For the exposure phase—which only measured scene recognition—discriminability (d’) was significantly greater for high- vs. low-training scenes (β = 0.473, t_(101)_ = 3.887, P < 0.001; **Fig. 3b**). For the post-test, associative memory accuracy was again better for high- vs. low-training scenes (β = 0.291, t_(101)_ = 9.320, P < 0.001; **Fig. 3c**), confirming that the benefits of high training (in the training phase) persisted throughout the experiment.

**Fig. 3.**
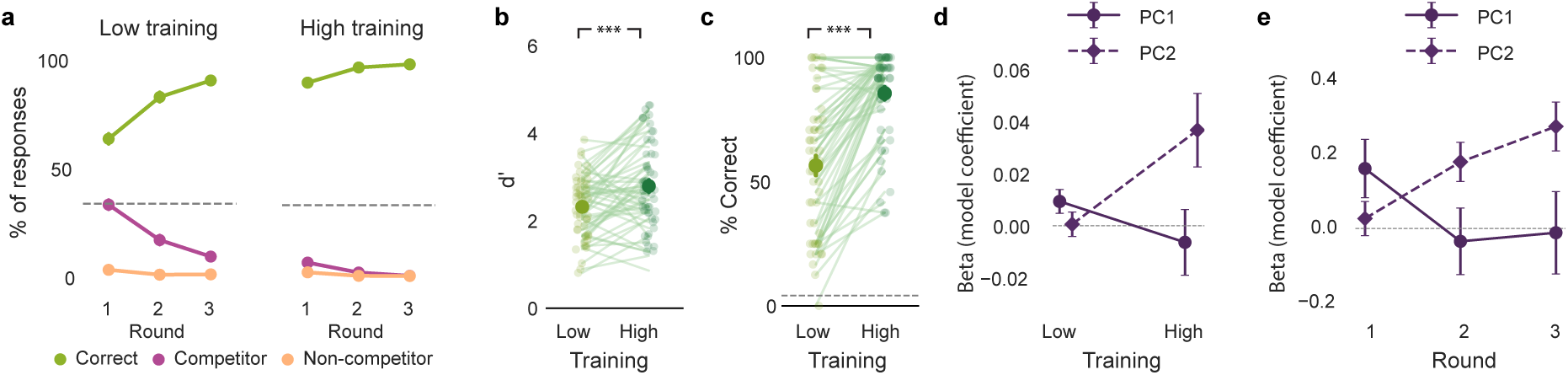
Influence of training on behavior. **a**, Performance during the training phase of the fMRI experiment. Left, associative memory accuracy (% correct responses) for low- vs. high-training scenes during each round. Circle markers represent mean accuracy and error bars indicate S.E.M. Right, percentage of responses corresponding to selection of the competitor vs. non-competitor, separately for low- vs. high-training scenes. **b**, Memory sensitivity (d’) during the exposure phase of the fMRI experiment for low- vs. high-training scenes. **c**, Associative memory accuracy during the post-test of the fMRI experiment for low- vs. high-training scenes. In **a** and **c**, dashed lines represent chance performance. For all bar plots in **a-c**, bars represent the mean, and the error bars indicate S.E.M. Asterisks reflect significant paired samples t-tests (***P < 0.001). **d**, The influence of each similarity dimension (PC1, PC2) on interference errors during the training phase of the fMRI experiment, separately for low vs. high training categories. To ensure individuals with few errors are not driving the effects in the high training condition, we repeated the analyses after excluding the participants with higher than the full sample’s mean performance. Even then, with high training, there was a significant main effect of PC2 degree centrality (*β* = 0.0398, t_(2076)_ = 2.560, P = 0.011) with no effect of PC1 degree centrality (*β* = −0.0035, t_(2076)_ = −0.258, P = 0.797). **e**, The influence of each similarity dimension (PC1, PC2) on interference errors during the training phase of the behavioral memory experiment, separately for each round of training (round 1 = low training, round 3 = high training). In plots **d** and **e**, the circle (for PC1) and diamond (for PC2) markers indicate beta estimates from linear mixed-effects models in which trial-by-trial degree centrality values (from PC1 and PC2) were used to predict the likelihood of an interference error (selecting the competitor). Error bars represent the corresponding S.E.M.s from the models.

We next tested whether the relative influence of each dimension of similarity on memory errors changed as a function of training. To test this, we focused on performance during the test rounds of the training phase (of the fMRI experiment). Specifically, we used mixed-effects models in which the dependent variable represented trial-by-trial interference errors (selecting the competitor; **Fig. 2a**). The key independent fixed effects were the degree centrality of the presented scene, separately for PC1 and PC2. We predicted that scenes with higher degree centrality (stronger ‘connections’ to other scenes; **Fig. 2d**) would be associated with more interference errors.

For PC2, there was a significant interaction between degree centrality and training (β = 0.036, t_(7464)_ = 2.408, P = 0.016; **Fig. 3d**); for PC1, the interaction was qualitatively opposite, but not significant (β = −0.015, t_(7464)_ = −1.110, P = 0.267; **Fig. 3d**). A linear contrast (F-test) confirmed a significant difference between these two effects (β = 0.051, F_(1, 7464)_ = 4.144, P = 0.042; **Fig. 3d**). Qualitatively, the influence of PC1 on memory errors was stronger with low training, whereas the influence of PC2 was stronger with high training. Follow-up tests (contrasting coefficients to a reference value of 0) confirmed that for the low-training condition there was a significant effect of PC1 (β = 0.0093, t_(3732)_ = 2.044, P = 0.041), but not PC2 (β = 0.0005, t_(3732)_ = 0.111, P = 0.912) whereas for the high-training condition there was a significant effect of PC2 (β = 0.0367, t_(3732)_ = 2.596, P = 0.009), but not PC1 (β = −0.0064, t_(3732)_ = −0.510, P = 0.610). For effects separated by training round, see **Supplementary Fig. 4**.

We next sought to conceptually replicate the preceding findings using data from the training phase of the behavioral memory experiment. Because the behavioral memory experiment did not manipulate training level (high vs. low), we instead focused on changes across the three training rounds, with the rationale being that early training rounds correspond to ‘low’ training and later training rounds correspond to ‘high’ training. As in the fMRI experiment, for PC2, there was a significant interaction between degree centrality and training (β = 0.0058, t_(17560)_ = 2.817, P = 0.005; **Fig. 3e**), and a qualitatively opposite effect for PC1 (β = −0.0027, t_(17560)_ = −1.654, P = 0.098; **Fig. 3e**). Again, these effects were significantly different from each other (β = 0.0085, F_(1, 17560)_ = 7.223, P = 0.007; **Fig. 3e**). With low training (here, round 1), there was a significant effect of PC1 (β = 0.0041, t_(5852)_ = 2.059, P = 0.040), but not PC2 (β = 0.0014, t_(5852)_ = 0.585, P = 0.559), and with high training (round 3), there was a significant effect of PC2 (β = 0.0146, t_(5852)_ = 4.171, P < 0.001), but not PC1 (β = −0.0003, t_(5852)_ = −0.102, P = 0.919). Thus, the pattern of statistical results was entirely consistent across the two experiments, despite minor differences in the paradigm and models. Collectively, these experiments demonstrate that both dimensions of similarity contributed to behavioral interference, but these influences occurred *sequentially* across training. Whereas PC1 drove errors at earlier stages of learning, PC2 drove errors at later stages of learning.

### Relationships between experience, similarity dimensions, and the hippocampus

We next turned to the central question of how visual similarity and experience influenced representational structure in the hippocampus. Based on prior studies in humans^15,27–32^ and rodents^1,4,5,33^, we had an *a priori* interest in hippocampal subfields CA3 and dentate gyrus (CA3/DG). For comparison, we also included subfield CA1 and two visual regions: early visual cortex (EVC) and parahippocampal place area (PPA). We anticipated that effects of visual similarity and experience would qualitatively differ between CA3/DG and the visual regions^16,17,27,28^. All fMRI analyses were based on measures of *relative* pattern similarity obtained during the exposure phase of the fMRI experiment. Specifically, for each participant and region, we computed the fMRI pattern similarity (Pearson correlation) for scenes from the same category (within-category similarity) and expressed this relative to similarity between scenes from different categories (across-category similarity) (**Fig. 4a**)^16,17,27,28^. We refer to the difference between these measures (within – across) as the *fMRI similarity score*.

**Fig. 4.**
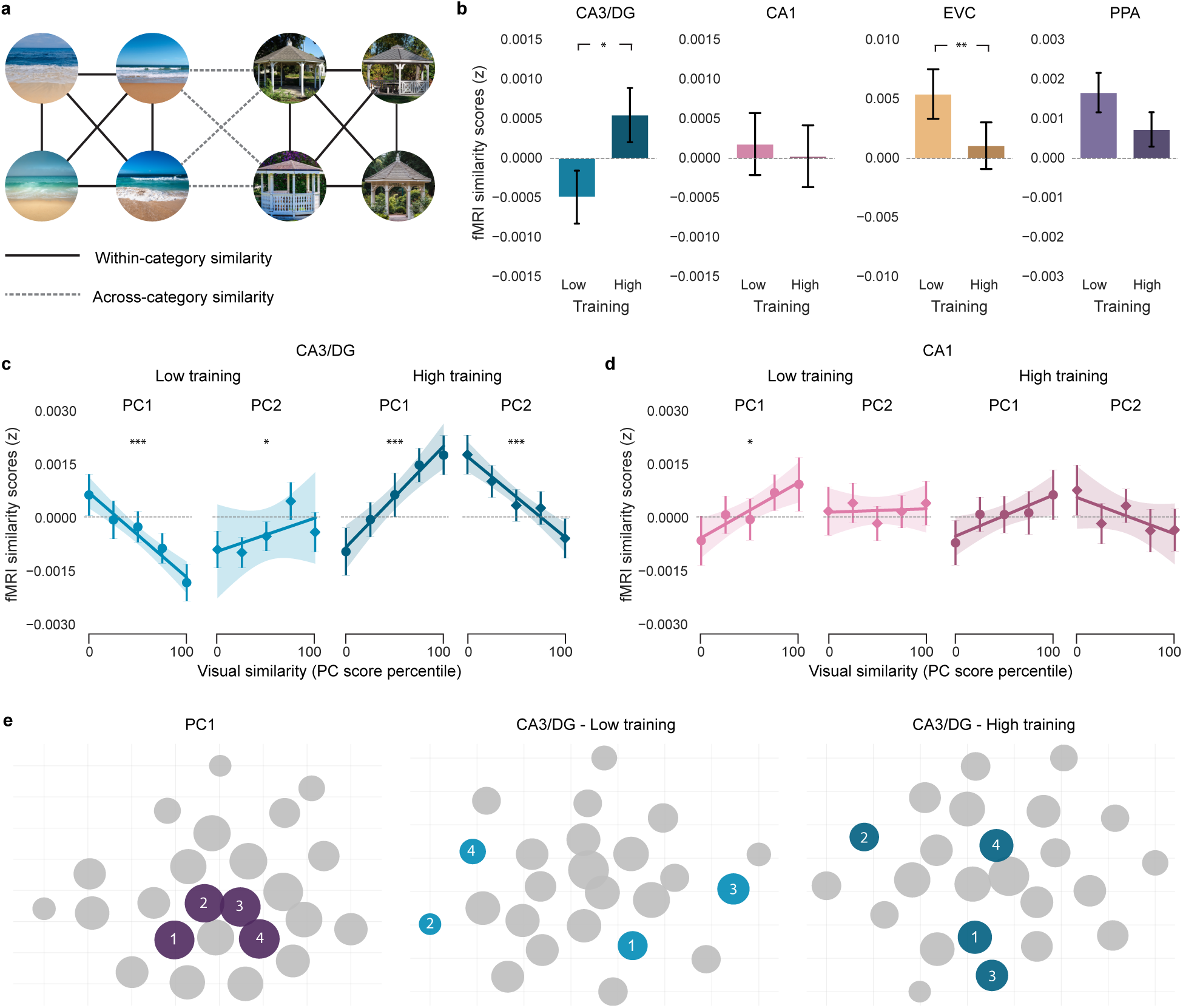
Relationships between similarity dimensions, experience, and fMRI pattern similarity. **a**, Schematic showing within- vs. across-category pattern similarity. **b**, Mean fMRI similarity scores by training level for CA3/DG, CA1, EVC, and PPA. Error bars reflect S.E.M. **c**,**d**, Relationships between visual similarity (PC1 and PC2 scores; circle vs. diamond markers, respectively) and similarity scores in CA3DG (**c**) and CA1 (**d**), separately for each training level (low, high). For **c,d**, statistical analyses were based on linear mixed effects models in which visual similarity (PC scores) were a continuous variable. Solid lines reflect lines of best fit and shaded area reflects S.E.M. Asterisks above plots indicate significant effects of visual similarity (PC score) on fMRI similarity scores (*P < 0.05, **P < 0.01, ***P < 0.001). Markers (circles, diamonds) reflect mean similarity scores at each of 5 levels (bins) of visual similarity. Error bars on markers reflect S.E.M. Binned data are purely for visualization purposes (as in Fig. 2c). **e**, 2-D visualization, using t-distributed Stochastic Neighbor Embedding (t-SNE), of the representational structure of the 24 scenes in the beach category, separately for visual similarity space (PC1, left), CA3/DG with low training (center) and CA3/DG with high training (right). Each circle reflects on scene. Circle size is proportional to a scene’s degree centrality in the associated space. Distance between circles is proportional to similarity. The four scenes with the highest degree centrality in the visual similarity space are shown in purple (left) and shown again in CA3/DG (light blue in the center and dark blue on the right).

As with our behavioral analyses, we first tested for overall effects of training—that is, whether fMRI similarity scores differed for high vs. low training (**Fig. 4b**). Linear mixed-effects models revealed that, within the visual regions, fMRI similarity scores decreased with training (EVC: β = −0.0044, t_(100)_ = −3.234, P = 0.002; PPA: β = −0.0009, t_(100)_ = −1.807, P = 0.074). With low training, similarity scores were positive (EVC: t_(51)_ = 2.594, P = 0.012, d = 0.360; PPA: t_(51)_ = 3.339, P = 0.002, d = 0.463) but did not differ from 0 with high training (|t_(51)_’s| < 1.688, P’s > 0.097, d’s < 0.234). In sharp contrast, similarity scores in CA3/DG significantly *increased* with training (β = 0.0010, t_(101)_ = 2.186, P = 0.031). Qualitatively, this reflected a transition from negative similarity scores (low training) to positive similarity scores (high training). However, similarity scores did not significantly differ from 0 at either training level (|t_(51)_’s| < 1.596, P’s > 0.116, d’s < 0.222). For CA1, there was no effect of training (β = 0.0002, t_(101)_ = −0.283, P = 0.777) and similarity scores did not differ from 0 at either training level (|t_(51)_’s| < 0.462, P’s > 0.646, d’s < 0.065). Thus, training influenced representational structure within visual cortical ROIs and CA3/DG, but in entirely opposite ways. It is notable that CA3/DG representations were less similar (greater *differentiation*) in the low training condition despite the fact that, overall, participants experienced more interference in the low training condition (**Fig. 3a**).

We next tested whether similarity within CA3/DG was related to the two dimensions of stimulus similarity (PC1, PC2) and whether these relationships depended on experience (training). While counter-intuitive, prior studies have shown that hippocampal representations are sometimes *less similar* when visual stimuli are *more similar*^15–17,27,28,30,34–39^. This phenomenon—referred to as ‘repulsion’—suggests that the hippocampus adaptively responds to memory interference by exaggerating differences between memories that are most similar (or most confusable). Thus, we predicted that the hippocampus (CA3/DG, in particular) would *invert* dimensions of similarity specifically when those dimensions were actively contributing to interference. Based on our behavioral findings (**Fig. 3d**), this translates to the low training condition for PC1 and the high training condition for PC2.

In contrast to the fMRI analyses described above, which computed the average similarity for all pairs of images within a scene category, here we were interested in fine-grained gradations in similarity within a category. Specifically, for each ROI, we ran linear mixed-effects models for which the dependent measure was fMRI pattern similarity for each of the 552 within-category image pairs (276 beach pairs + 276 gazebo pairs). Fixed effects included PC1 and PC2 scores (continuous predictors included in the same model) and training level (low vs. high).

For CA3/DG, the interaction between PC score and training level was highly significant for both PC1 and PC2, but these interactions took opposite forms (PC1: β = 0.0018, t_(28697)_ = 8.628, P < 0.001; PC2: β = −0.0013, t_(28697)_ = -6.287, P < 0.001; **Fig. 4c**). Specifically, with low-training, PC1 scores were *negatively* related to CA3/DG similarity (β = −0.00085, t_(14348)_ = −4.656, P < 0.001) while PC2 scores were *positively* related to CA3/DG similarity (β = 0.00047, t_(14348)_ = 1.981, P = 0.048). With high training, these relationships *flipped*: (PC1: β = 0.00100, t_(14348)_ = 3.910, P < 0.001; PC2: β = −0.00089, t_(14348)_ = - 3.408, P < 0.001). In other words, with low training, CA3/DG inverted similarity along PC1, but preserved similarity along PC2; with high training, CA3/DG inverted similarity along PC2, but preserved similarity along PC1. The fact that CA3/DG selectively inverted PC1 during low training, and PC2 during high training strikingly parallels our behavioral findings that PC1 drove interference during low training and PC2 drove interference during high training. In other words, CA3/DG selectively inverted each dimension of similarity when it was contributing to memory interference. These transformations were putatively adaptive because they dramatically—but transiently—increased the representational distance between the pairs of images that were most at risk for interference (**Fig. 4e**).

As noted earlier, our interpretation of PC1 and PC2 is that these dimensions roughly captured perceptual and semantic similarity, respectively. To reinforce this interpretation, we implemented a mixed-effects model, identical to the preceding model, but instead of using PC1 and PC2, we selected a single metric that putatively captures perceptual similarity (VGG MP1) and a single metric that putatively captures semantic similarity (NLP of text descriptions). Qualitatively, we observed an identical pattern of data (**Supplementary Fig. 5**). Namely, with low training CA3/DG inverted perceptual similarity (P = 0.002); with high training, CA3/DG inverted semantic similarity (P = 0.008). For more detailed consideration of how individual metrics related to CA3/DG similarity, see **Supplementary Fig. 6, 7 and 8**. Notably, we also found that PC1 and PC2 were highly robust to the removal of individual similarity metrics (this was particularly true for PC1; see **Supplementary Table 2**).

To further support the argument that repulsion within CA3/DG was a direct reaction to memory interference, we implemented another mixed-effects model, this time using similarity values taken directly from the two behavioral similarity experiments: the behavioral memory experiment and the perceptual similarity experiment (**Fig. 1c**). Strikingly, across training levels, CA3/DG similarity was negatively related to similarity derived from the memory experiment (β = −0.00062, t_(28692)_ = -2.157, P = 0.031), with a qualitatively opposite effect for the perceptual experiment (β = 0.00069, t_(28692)_ = 1.390, P = 0.165; **Supplementary Fig. 9**). In other words, when fully controlling for perceptual similarity judgments, CA3/DG was sensitive to—and *inverted*—dimensions of similarity that contributed to memory interference.

Interestingly, independent of any metrics or dimensions of similarity, we also observed that, with high training, CA3/DG inverted the representational structure that was present with low training (**Supplementary Fig. 10**), further demonstrating that CA3/DG transformed information across stages of learning.

Notably, the relationships between PC1/PC2 similarity and training that we observed in CA3/DG were not evident in CA1 (**Fig. 4d**) or in the visual regions (**Supplementary Fig. 11**). Additionally, training-related effects in CA3/DG were absent for other PCs (PCs 3-10; **Supplementary Table 3**).

## Discussion

Relationships between visual similarity and hippocampal similarity are at the core of theories of hippocampal function^1–5^. Yet, these relationships can take qualitatively different forms^5,12,14–17,27,28,30,31,34–41^. Here, we show that the shape of input-output functions critically depends on the current relevance of individual dimensions of visual similarity to behavioral expressions of memory interference. We first identified two dimensions of visual similarity that robustly contributed to memory interference. We then show that these dimensions of similarity influenced behavior *sequentially*, across stages of learning. Finally, in CA3 and dentate gyrus (CA3/DG), we reveal a striking parallel to these behavioral findings: across learning, CA3/DG inverted each dimension of similarity precisely when it contributed to memory interference. These findings provide important insight into when and why the hippocampus transforms dimensions of visual similarity. Specifically, hippocampal input-output functions dramatically change with experience, and these changes reflect a dynamic prioritization of dimensions of similarity that are most relevant to behavior.

The major innovation in our approach is that we rigorously characterized different dimensions of visual similarity. We did this by aggregating 10 similarity metrics and using PCA to extract orthogonal components (dimensions). Thus, rather than operationalizing dimensions based on an assumed 1-to-1 mapping (metric X = dimension Y), our approach allowed dimensions to be *distributed* across metrics (**Fig. 1i**). Importantly, the dimensions we identified were behaviorally relevant—participants were much more likely to confuse pairs of images that had high PC1 or PC2 scores (**Fig. 2c**). While many prior behavioral studies also show that similarity drives memory interference^7–11^ and that interference occurs along specific visual dimensions^42^, our findings reveal an important caveat: different dimensions of similarity drive interference at different stages of learning. Indeed, we show—and replicate—that during early stages of learning, there was an influence from PC1, *but not PC2*; in late stages of learning, there was an influence from PC2, *but not PC1* (**Fig. 3d-e**).

The fact that dimensions of similarity drove interference in serial (not parallel), is reminiscent of—and consistent with—findings from the perceptual discrimination literature which show that, when categorizing multidimensional stimuli, humans favor unidimensional strategies^43–47^. Here, PC1 putatively drove interference errors first (followed by PC2) because, by definition, PC1 explained more variance in the similarity matrices (41.3%) than PC2 (19.1%)^46^. The fact that PC1 had no influence during later stages of learning indicates that interference on this dimension was successfully resolved. While caution is warranted when explicitly labeling or describing PCA components, the observed loadings for PC1 and PC2 (**Fig. 1i**) suggest that PC1 reflected relatively lower-level/perceptual information whereas PC2 reflected higher-level/semantic information. This interpretation is potentially compatible with other evidence showing that, with experience, memories transition from perceptual to semantic information^48–51^. That said, we do not predict that the relative prioritization of PC1 vs. PC2 observed here will necessarily generalize across other stimulus sets and procedures. Rather, we believe the current findings reflect a generalizable *principle*: that interference is resolved ‘one dimension at a time’. For example, by changing task demands or the composition of the stimulus set, it may be possible to manipulate which dimensions of similarity are relevant at each stage of learning. However, the key prediction would be that even if dimensions are re-prioritized, interference would still be resolved one dimension at a time.

By considering different dimensions of visual similarity and different levels of experience, we show that relationships between visual similarity and hippocampal similarity (input-output functions) can take remarkably different forms, ranging from strongly positive to strongly negative (**Fig. 4c**; **Supplementary Table 3**). These findings add to recent evidence that relationships between visual similarity and hippocampal similarity are experience-dependent^17,27,28^. However, our behavioral findings provide a compelling framework for understanding these experience-dependent changes.

Namely, for PC1 and PC2, CA3/DG either preserved or inverted each dimension according to whether it *actively* contributed to interference. Thus, CA3/DG had access to diverse visual input^52^, but *selectively* transformed this input when it was adaptive for behavior^53–55^.

The fact that we specifically implicate CA3/DG in resolving memory interference (as opposed to CA1) is consistent with prior human and rodent studies^15,31,36,37,56–58^. In particular, CA3/DG has been implicated in *orthogonalizing* similar memories (pattern separation)^1,5,14^. That said, standard accounts of pattern separation do not predict or explain negative input-output functions^1–5^. In fact, with perfect orthogonalization, input-output functions would be flat, not negative (for additional consideration of this point, see^27,59^). Thus, while reinforcing the relevance of CA3/DG to memory interference, our findings also challenge existing models of how CA3/DG resolves interference.

An important aspect of our design and analyses is that we densely sampled a narrow range at the high end of similarity space. Namely, our models included 276 pairwise comparisons within each visual category. In contrast, the vast majority of prior empirical evidence for negative input-output functions in the hippocampus comes from studies that have only sparsely sampled similarity space^15–17,27,28,30,34,35,37,39,60^ (high vs. low similarity; *cf*.^34^). More broadly, the input-output functions we observed in CA3/DG (positive and negative) indicate a striking degree of sensitivity to gradations within a relatively narrow similarity space. In fact, CA3/DG was more sensitive to within-category differences (e.g., differences between beaches) than across-category differences (differences between beaches and gazebos) (i.e., within-category similarity did not systematically differ from across-category similarity; **Fig 4b**).

Although we primarily focus on input-output functions within each visual category, we also observed a surprising main effect of training in CA3/DG: lower similarity scores (greater differentiation) in the low training condition compared to the high training condition (**Fig. 4b**). This effect strongly contrasted with EVC, which showed a robust *decrease* in similarity scores with training. While speculative, these opposing effects are potentially interrelated: CA3/DG differentiation may be most likely to occur when input from visual cortical areas is un-differentiated^16,17,27,28^. From this perspective, if visual cortical areas differentiate stimuli (through experience-dependent attention to diagnostic features^61–63^), then there is less need for additional differentiation by CA3/DG^27,38^.

In summary, we show that, in behavior and the hippocampus, memory interference is resolved one dimension at a time. When a particular dimension of visual similarity is actively contributing to memory interference, CA3 and dentate gyrus dramatically—but transiently—invert that dimension. These findings can be interpreted through a simple framework in which the hippocampus selectively exaggerates the representational distance between memories that are most at risk for interference.

## METHODS

### Participants

Human subject data were collected for three experiments: the behavioral memory experiment, the perceptual similarity experiment, and the fMRI experiment. Across the three experiments, a total of 655 participants were enrolled. All participants’ enrollment followed procedures approved by the University of Oregon Institutional Review Board. For the behavioral memory experiment and the fMRI experiment, all participants were right-handed native-English speakers. For the fMRI experiment, all participants had normal or corrected-to-normal vision, with no self-reported psychiatric or neurological disease.

For the behavioral memory experiment, data were collected from 463 participants (200 female, mean age = 28.02, range = 18 – 35 years) using the online platform Prolific. Informed consent was obtained, online, for each participant prior to the experiment. To ensure that the memory confusability matrix obtained from this experiment was highly reliable, participants with low performance were excluded (see ‘Experimental procedure – Behavioral memory experiment’ and **Supplementary Fig. 12**). The final analysis included data from 122 participants. Participants in the behavioral memory experiment received monetary compensation.

For the perceptual similarity experiment, data were collected from 141 undergraduate students from the University of Oregon Human Subjects pool (95 female, mean age = 19.04, range = 18 – 27 years) that completed the experiment online through the Sona system. Informed consent was obtained, online, for each participant prior to the experiment. There were two versions of the perceptual similarity experiment: one that only included images of beaches, and another with images of gazebos. Of the 141 participants, 71 participated in the beach experiment and 70 participated in the gazebo experiment. To ensure that the perceptual similarity measure from this experiment was reliable, participants with an insufficient amount of data and/or low performance were excluded (see ‘Experimental procedure – Perceptual similarity experiment’). The final analysis included data from 113 participants—57 for the beach experiment and 56 for the gazebo experiment. Participants in the perceptual similarity experiment were compensated with study credit.

For the fMRI experiment, 53 participants (42 female, mean age = 21.23, range = 18 – 31 years) completed the experiment with a target sample of at least 50. Because relevant effect sizes were not available (for the effects of interest here), we did not perform a formal sample size calculation. Rather, the goal was simply for the experiment to be well powered. Written informed consent was collected for each participant prior to the experiment. One participant’s data was excluded due to a technical error in the MRI acquisition. All analyses of fMRI data are based on 52 participants. Participants in the fMRI experiment received monetary compensation.

### Stimuli

All three experiments used the same set of 24 images of beaches and 24 images of gazebos (**Fig. 1a**). For the fMRI experiment, a separate set of 24 gazebos and 24 beaches were used as lures for the exposure phase. All scene images were color photographs, drawn from Google Images. The behavioral memory experiment and the fMRI experiment also used 48 images of everyday objects (color photographs, drawn from Google Images) shown against a white background. Object images were selected such that no two objects shared a common label; otherwise, similarity among objects was not measured or controlled. All scene images and objects were square, available in the public domain, and will be shared by the time of publication at https://osf.io/gy23h/ as part of the manuscript’s dataset. For the fMRI and behavioral memory experiments, the 48 scenes were randomly paired with the 48 objects, separately for each participant, to form 48 scene-object associations. For the fMRI experiment, one scene category (e.g., gazebos) was assigned as the ‘high-training’ category, and the other scene category (e.g., beaches) was assigned as the ‘low-training’ category. The high- and low-training category assignment alternated across participants.

### Experimental procedure

#### Behavioral memory experiment

The behavioral memory experiment, conducted online (using PsychoPy’s online platform, Pavlovia), involved learning scene-object associations. It consisted of a training phase followed by a post-test. Across all phases, stimuli were presented on a gray background on a computer screen. The experiment was implemented in PsychoPy2024.1.5 and lasted for about 35 minutes. The training phase consisted of 3 study-test rounds and was most similar to the low training condition of the fMRI experiment (described below). In each study/test round, all 48 scene-object associations were studied/tested once, in random order. The assignment of objects to scenes was random for each participant but was stable throughout the experiment. For each trial of the study rounds, a scene and its associated object were displayed next to each other on the screen for 1500 ms (scene on left, object on right), with a white fixation cross in between. Study trials were separated by a 1000 ms inter-trial interval during which a fixation cross was displayed. For each trial of the test rounds, a scene (probe) was displayed at the top of the screen, with 3 objects presented underneath. One of the objects was the target (the object that was paired with the probe during the study rounds). One object was the competitor (an object that was paired with a scene from the same category as the probe). One object was the non-competitor (an object that was paired with a scene from the opposite category as the probe). In each test round, each object appeared once as the target, once as the competitor, and once as the non-competitor. For each scene, the competitor and non-competitor objects were randomly and independently selected in each round. Participants had 10 s to select one of the objects using the keyboard. After making a response, or after 10 s elapsed, the trial ended. Test trials were separated by a 1000 ms inter-trial interval during which a fixation cross was displayed.

The post-test consisted of three rounds, during which each of the 48 studied scenes was tested once per round, in random order. On each post-test trial, a scene (probe) was displayed at the top of the screen, with 24 objects displayed below (3 rows x 8 columns). One of the 24 objects (target) had been paired with the probe. The other 23 objects (competitors) had been paired with a scene from the same category as the probe. Participants were instructed to use their trackpad or mouse to select the object that had been paired with the scene. Participants had 30 seconds to respond on each trial. After responding, or after 30 s elapsed, the trial ended. Post-test trials were separated by a 500 ms inter-trial interval during which a fixation cross was displayed. Post-test data were used to generate a confusability matrix (see ‘Metrics of Similarity,’ below). The rationale for testing each scene three times during the post-test was to obtain a more reliable confusability matrix (for validation, see **Supplementary Fig. 1c**).

Because the behavioral memory experiment was used to establish the normative confusability of scene pairs, it was important that the data were internally reliable. Thus, separately for each scene category, we excluded participants whose mean accuracy across the three post-test rounds was below 28%. While this threshold was well above chance (4.17%), we selected this threshold to optimize split-half reliability of the memory confusability measure.

#### Perceptual similarity experiment

The perceptual similarity experiment, conducted online (using PsychoPy’s online platform, Pavlovia), involved making subjective judgments about the relative similarity of scene images. It was modeled after the triplet odd-one-out task described in^20^. Stimuli were presented on a gray background on a computer screen. The experiment was implemented in PsychoPy2024.1.4 and each session lasted approximately 15 minutes. In short, participants completed a single task in which three scenes were shown, and participants select the ‘odd one out’ (the scene that was least similar to the other two scenes). Each participant in the experiment made perceptual similarity judgments for one of the two visual categories (beach experiment or gazebo experiment). For each experiment (beach or gazebo), participants were allowed to enroll in a second session (for the same category) to be completed within three days of the first session for additional credit. The second session presented triplets that were not shown in the first session. A minimum two-hour break between sessions was enforced. For each category, because there were 24 scenes, there were 2,024 possible triplets. For each participant, the sample of triplets was randomly and independently chosen. On each trial, one of the 2,024 triplets was randomly selected (without replacement within participants). The corresponding three images were presented equidistantly on a virtual circle. On each trial, the image array was randomly rotated by 1–360 degrees to prevent predictable locations and discourage habitual responses. This prevented participants from repeatedly selecting the same screen location without evaluating the presented options. Participants selected the odd image by clicking on it with the trackpad or mouse. There was no time limit for responses. Trials were separated by a 500 ms inter-trial interval during which a fixation cross was displayed. Each session lasted 15 minutes with a self-terminated break after every 50 trials. On average, each participant completed a total of 339 trials per session (SD = 106.0) in the beach experiment and 314 trials per session (SD = 73.4) in the gazebo experiment. 5% of trials were ‘catch trials’ in which two of the scenes were identical. The catch trials were used to identify participants responding randomly. Participants were not informed about the presence of catch trials. Participants were excluded if their accuracy on catch trials (selecting the objectively different image) was lower than 80%. Data from the perceptual similarity experiment were used to generate a perceptual similarity matrix (see ‘Metrics of Similarity,’ below). The 80% threshold for exclusion was chosen to optimize split-half reliability of the perceptual similarity matrix.

#### fMRI experiment

The fMRI experiment, which was entirely in-person, was similar to the behavioral memory experiment, with the two biggest differences being (1) during the training phase, one category received ‘high training’ and the other category received ‘low training,’ and (2) the training phase and the post-test were separated by an exposure phase. As in the behavioral memory experiment, objects were randomly paired with scenes for each participant, but these associations were stable throughout the experiment. The training phase was conducted in a preparation room, using a laptop, prior to entering the MRI scanner. Inside the scanner, participants completed eight rounds (functional scans) of the exposure phase. Participants then exited the scanner and completed the post-test, using a laptop, in the preparation room. Across all phases, stimuli were presented on a gray background, either directly on a laptop screen or projected from the back of the MRI scanner. The experiment was implemented in PsychoPy2022.1.1 and lasted for about 2 hours and 30 minutes, with about 1 hour and 15 minutes inside the MRI scanner.

The training phase consisted of 3 study-test rounds. In each study/test round, all 48 scene-object associations were studied/tested in random order. Associations in the high training condition were studied three times per round and tested once per round. Associations in the low training condition were studied/tested once per round. For each trial of the study rounds, a scene and its associated object were displayed next to each other on the screen (scene on left, object on right), with a white fixation cross in between. The display time for each study trial varied by training condition (high vs. low) and study/test round. For the high-training category, the display times were: 3000 ms (round 1), 2500 ms (round 2), and 2000 ms (round 3). For the low-training category, the display times were: 2500 ms (round 1), 2000 ms (round 2), and 2000 ms (round 3). Study trials were separated by a 500 ms inter-trial-interval during which a fixation cross was displayed. For each trial of the test rounds, a scene (probe) was displayed at the top of the screen, with 3 objects presented underneath. As in the behavioral memory experiment, one of the objects was the target, one was the competitor, and one was the non-competitor. In each test round, each object appeared once as the target, once as the competitor, and once as the non-competitor. For each scene, the competitor and non-competitor objects were randomly and independently selected in each round. Participants had 10 s to select one of the objects using keyboard. For the high-training category, immediately after a response was made, or after 10 s elapsed, participants received feedback: the two incorrect objects were removed and only the correct object remained on the screen (1000 ms). Feedback was not provided for the low-training category. Test trials were separated by a 1000 ms inter-trial interval during which a fixation cross was displayed.

The exposure phase was conducted during fMRI scanning. In each of the 8 rounds (scan runs), participants were shown each of the 48 previously-studied scenes (24 beaches, 24 gazebos) along with 6 new ‘lure’ scenes (3 beaches, 3 gazebos). The 54 scenes were presented one at a time, in random order. The lures never repeated across scan runs. Thus, the exposure phase contained a total of 6 (lures) * 8 (runs) = 48 lures. Participants’ only task during the exposure phase was to make a button press using their index finger when they detected a new scene—i.e., one not presented during the study phase. Performance was evaluated in a signal detection framework, with button presses for objectively new scenes treated as ‘hits’ and button presses for objectively old scenes treated as ‘false alarms.’ On each trial, a scene was displayed for 2000 ms, followed by 4000 ms inter-trial interval during which a white fixation cross was displayed. Participants were instructed to respond (if relevant) by the end of the fixation period. Participants were not instructed or encouraged to practice recalling the object associations during the exposure phase and they were not made aware that scene-object memory would be tested again in a post-test. By design, the exposure phase had minimal behavioral demands to resolve interference. The rationale for this was to minimize potentially confounding effects of behavioral responses/decisions. Instead, the exposure phase was intended to capture consequences of prior learning (training) and stimulus similarity.

The post-test consisted of a single round in which each of the 48 studied scenes was tested once, in random order. On each post-test trial, a scene (probe) was displayed at the top of the screen, with 24 objects displayed below (3 rows x 8 columns). As in the behavioral memory experiment, one of the 24 objects was the target, and the other 23 objects were competitors. Participants were instructed to use the trackpad to select the object that had been paired with the scene. Participants had 99 seconds to respond on each trial. After responding, or after 99 s elapsed, the trial ended. Post-test trials were separated by a 500 ms inter-trial interval during which a fixation cross was displayed.

### Metrics of similarity

We selected 10 different metrics of stimulus similarity. While the selection of metrics was based on subjective judgment of the experiments (as opposed to objective criteria), the intention was to combine human-based judgments and computational methods. For the human-based judgments, our intuition was that subjective judgments of perceptual similarity may be at least partially distinct from direct measures of memory confusability. For the computational methods, our goal was to span low-, mid-, and high-level perceptual similarity as well as more conceptual/semantic similarity. The 10 metrics are described below.

#### Memory confusability

Responses from the post-test of the behavioral memory experiment were used to generate a normative memory confusability matrix. After excluding participants based on performance thresholds (see ‘Experimental procedure – Behavioral memory experiment’), a matrix was generated for each participant that reflected the pairwise confusability for each pair of scenes within a category. For example, in a post-test round, when presented with beach 3 during the post-test, if participants selected the object that had been paired with beach 7 (see **Fig. 1c**, left), this would be considered a ‘confusion’ between beach 3 and beach 7 (coded as ‘1’ in the confusability matrix for that round, see **Fig. 2b**). For each participant, the confusion matrices over the 3 post-test rounds were averaged, constituting the confusion matrix for that participant. For example, if beach 3 was confused with beach 7 in 2 out of the 3 post-test rounds, that element of the confusability matrix would be 0.667. Participant-level matrices were then averaged. Because the resulting distribution of confusability values was skewed (many pairs were rarely confused, resulting in a left-skew), a log transformation was applied, yielding a distribution of confusability values that was approximately normal.

#### Perceptual similarity

Responses from the perceptual similarity experiment were used to generate a normative perceptual similarity matrix. Similarity for a given pair of scenes (e.g., beach 2, beach 7 similarity) was defined as the proportion of trials, when beach 2 and beach 7 were presented together, on which the other scene (the third scene) was selected as the ‘odd one out’. In other words, the proportion of trials when the ‘other’ scene was selected reflects the similarity of a given pair of scenes relative to other possible pairs of scenes. For each participant, a similarity matrix was generated where each cell represented the proportion of ‘other’ responses. After performance-based exclusion of participants (see the details in ‘Experimental procedure – Perceptual similarity experiment’), similarity matrices were averaged across participants. Because the resulting distribution of similarity values was skewed (many pairs had low/zero similarity, resulting in a left-skew), a log transformation was applied, yielding a distribution of similarity values that was approximately normal.

#### CLIP

To quantify the similarity between scenes using CLIP^64^, the scene images were passed through the image encoder of the CLIP model (ViT-B/32 variant), a transformer-based deep learning architecture pretrained on a large corpus of paired image-text data. The model and preprocessing pipeline were accessed using the open-source CLIP library (https://github.com/openai/CLIP) implemented in PyTorch. Each image was preprocessed according to the model’s specifications, encoded into a high-dimensional feature embedding, and normalized to unit length. Cosine similarity was then computed between all pairs of image embeddings for each scene category. CLIP captures visual and semantic relationships between images due to its joint image–text training.

#### NLP of text

To compute the similarity between scenes’ text descriptions using natural language processing (NLP), descriptive captions were first generated for each scene using OpenAI’s ChatGPT (GPT-4-turbo; gpt-4o variant), a multimodal large language model accessed through a ChatGPT Plus subscription. The prompt for each scene image was the following: ‘Please describe the visual content of this scene in one clear sentence, using no more than 20 words’. The resulting descriptions for each scene are shown in **Supplementary Fig. 2**. To compute pairwise similarity between the resulting text descriptions, embeddings were generated using the all-mpnet-base-v2 model from the Sentence-Transformers library. This model is based on MPNET, a deep learning architecture that integrates transformer-based attention with permutation-based pretraining to capture rich semantic information in text^65^. Sentence embeddings were computed for each description, and cosine similarity was calculated between each pair of scenes within each category. This approach primarily captured semantic similarity between scenes.

#### VGG-16

To quantify the similarity between scenes according to VGG-16^66^, we first extracted feature representations from the max pooling layers of a pretrained VGG-16 model using PyTorch (version 2.6.0). Input scene images were resized to 224×224 pixels and normalized according to VGG-16’s requirements. We passed each scene image through the convolutional feature extractor and collected activation maps from all five max pooling layers, though subsequent analyses focused specifically on layers 1, 3, and 5. Layers 2 and 4 were not considered given their high similarity to layers 1 and 3 respectively. Feature maps were flattened into one-dimensional vectors, and pairwise similarity between scenes was computed as the Pearson correlation between their corresponding feature vectors. This procedure yielded a layer-specific similarity matrix for each selected max pooling layer, reflecting progressively more abstract visual similarity as depth increased within the network hierarchy.

#### AlexNet

To measure the similarity between scenes according to AlexNet^67^, we used a pretrained AlexNet model using PyTorch (version 2.6.0). Each scene image was resized to 224×224 pixels and normalized using ImageNet preprocessing parameters before being passed through the network. 4096-dimensional feature vectors were extracted from the penultimate fully connected layer (fc7) using a forward hook. Pairwise similarity between scenes was computed using cosine similarity. Overall, this approach captures image-based similarity between scenes derived from the visual representations learned by the network.

#### GIST

GIST descriptor similarities between all pairs of scene images were computed using MATLAB R2023b’s LMgist implementation. Each scene image was converted to grayscale and resized to 256×256 pixels. GIST descriptors were calculated using 4 spatial scales with 8 orientations per scale, distributed across a 4×4 grid with a pre-filtering coefficient of 4. Similarity between scene image pairs was quantified as the cosine similarity between their GIST descriptor vectors. GIST similarity reflects the coarse structural and layout information of each scene, capturing its overall gist while ignoring fine-grained features^68^.

#### SSIM

Structural similarity indices (SSIM) were computed between all scene image pairs using MATLAB R2023b’s ssim function. Scene Images were converted to grayscale prior to analysis. The SSIM algorithm compared luminance, contrast, and structure between local image patches using an 11×11 Gaussian window, producing similarity values ranging from 0 (no similarity) to 1 (identical images) for each scene pair. SSIM quantifies how similar images are in terms of local luminance, contrast, and structure, emphasizing detailed perceptual features rather than global layout^69^.

### Dimensions of Similarity

For each of the 10 similarity metrics described above, we computed pairwise similarity for all 276 pairs of beaches and 276 pairs of gazebos, yielding 552 similarity values in total. We then constructed a 552 × 10 matrix, where each row corresponded to a scene pair and each column to a similarity metric. Principal component analysis (PCA) was applied to this matrix, yielding 10 orthogonal dimensions of similarity, with each scene pair assigned a score on each dimension (PC score).

Our rationale for applying PCA to the set of 10 similarity matrices (described above) was that different metrics are likely to capture shared variance. As such, similarity may be better explained by a few underlying dimensions than by individual metrics. We primarily focus on the first two dimensions (PC1 and PC2) because (1) by definition, they explained the most variance and (2) they strongly predicted memory interference (**Fig. 2c**). Importantly, PC1 and PC2 were each characterized by positive loadings from several metrics, with the loadings for other metrics closer to 0 (**Fig. 1i**). Based on the specific metrics that loaded positively on each PC, we interpreted them as reflecting perceptual similarity (PC1) and semantic similarity (PC2). To validate this interpretation, we submitted the PC loadings to ChatGPT using the following prompt:

‘Here are the loadings from the10 metrics of similarity. Please tell me what these two PCs of similarity capture in just one word/term.’

ChatGPT’s response was:

‘I would characterize them roughly as: PC1: Perceptual-structural similarity PC2: Semantic-conceptual similarity or even shorter: PC1: Perceptual PC2: Semantic.’

Thus, although our approach to identifying dimensions of similarity was entirely data driven (i.e., we did not impose *a priori* predictions), the resulting dimensions (PC1, PC2) are generally consistent with perceptual vs. semantic similarity, respectively. It is important to note that other dimensions (e.g., PC4), were characterized by a mix of positive and negative loadings (see **Supplementary Table 1**).

### MRI acquisition

All images were obtained on a Siemens 3T Skyra MRI system in the Lewis Center for Neuroimaging at the University of Oregon. Functional data were obtained using a T2*-weighted echo-planar imaging (EPI) sequence with partial brain coverage that prioritized full coverage of the hippocampus as well as visual cortex (repetition time = 2000 ms, echo time = 34 ms, flip angle = 90°, 66 slices, 1.7 × 1.7 × 1.7 mm voxels). Slices for functional scans were oriented parallel to the long axis of the hippocampus. A total of 8 functional scans were conducted. Each functional scan consisted of 168 volumes and included 6 s of lead-in time at the beginning and 6 s of lead-out time at the end of each scan. Anatomical scans included a whole-brain high-resolution T1- weighted magnetization prepared rapid acquisition gradient-echo anatomical volume (1 95 × 1 × 1 mm voxels) as well as a high-resolution (coronal direction) T2-weighted scan (0.43 × 0.43 × 1.8 mm voxels) to facilitate segmentation of hippocampal subfields.

### Anatomical data preprocessing

Preprocessing was conducted with fMRIPrep 23.2.0^70^ (RRID:SCR_016216), which is based on Nipype 1.8.6^71^ (RRID:SCR_002502). The T1-weighted (T1w) image was corrected for intensity nonuniformity (INU) with N4BiasFieldCorrection^72^ (ANTs 2.5.0^73^, RRID: SCR_004757) and used as the T1w reference throughout the workflow. The T1w reference was skull-stripped with the antsBrainExtraction.sh workflow (ANTs) in Nipype, using OASIS30ANTs as the target template. Brain tissue segmentation of cerebrospinal fluid (CSF), gray-matter (GM), and white-matter (WM) was conducted on the brain-extracted T1w using FAST^74^ (FSL; RRID:SCR_002823). The T2-weighted image was used to enhance pial surface refinement. Brain surfaces were reconstructed using recon-all (FreeSurfer 7.3.2, RRID:SCR_001847). Volume-based spatial normalization to one standard space (MNI152NLin2009cAsym) was conducted through nonlinear registration with antsRegistration (ANTs 2.5.0), using brain-extracted versions of T1w reference as well as the T1w template. ICBM 152 Nonlinear Asymmetrical template version 2009c^75^ was used for spatial normalization (RRID:SCR_008796; TemplateFlow ID: MNI152NLin2009cAsym).

### Functional data preprocessing

For each of the 8 functional (BOLD) scans per participant, preprocessing began with the generation of a reference volume for head motion correction. Head-motion parameters relative to this BOLD reference (six translation and rotation parameters plus the corresponding transformation matrices) were estimated. A fieldmap was collected at the scanner to correct for susceptibility-induced distortions. In addition, short EPI reference images with opposite phase-encoding directions were acquired; these were used in FSL’s topup to estimate distortions. If the first acquisition of the EPI references was of insufficient quality, it was repeated, resulting in two or more images. The raw fieldmap and EPI references were combined to produce a participant-specific distortion map (‘estimated fieldmap’). The estimated fieldmap was then rigidly aligned and applied to the reference EPI to correct distortions. The BOLD reference was next co-registered to the T1w reference using bbregister (FreeSurfer) which implements boundary-based registration^76^. Co-registration was configured with six degrees of freedom. Multiple time-series of potentially confounding variables were computed based on the preprocessed BOLD: framewise displacement (FD), DVARS, and three region-wise global signals^77^. Furthermore, a set of physiological regressors were extracted for component-based noise correction (CompCor)^78^. Principal components were estimated after high-pass filtering the preprocessed BOLD time-series (using a discrete cosine filter with 128 s cut-off) for the two CompCor variants: temporal (tCompCor) and anatomical (aCompCor). Frames exceeding a threshold of 0.5 mm FD or 1.5 standardized DVARS were annotated as motion outliers and excluded from the Generalized linear models (GLMs). The BOLD time-series were resampled onto fsaverage6. Gridded (volumetric) resampling was performed using nitransforms, configured with cubic B-spline interpolation.

Eight brain masks were generated using fMRIPrep for each of the eight functional scans per participant and the intersection of all eight masks was used as the final brain mask. The processed BOLD time-series were scaled to a mean of 100, with values clipped between 0 and 200. For the eight functional scans, all first-level GLMs were performed in participants’ native space with FSL using a Double-Gamma HRF with temporal derivatives, implemented in Python3.10. GLMs were calculated with AFNI’s 3dREMLfit^79^ using the Least Squares-Separate method^80^: for each scan, a separate GLM was calculated for each of the 48 scene images (24 beaches and 24 gazebos). Thus, for each GLM, there was one regressor of interest (representing a single scene image per scan). All other trials (including lure scene images), FD, xyz translation, xyz rotation, aCompCor 00-05, and csf were represented with nuisance regressors. This model resulted in 48 beta-maps per scan (one map per scene image) which were later converted to t-stats maps that represented the pattern of activity elicited by each scene in each functional scan.

### Regions of interest

A region of interest (ROI) for early visual cortex (EVC) was created from the probabilistic maps of Visual Topography^81^ in MNI space with a 0.5 threshold. This ROI was transformed into each participant’s native space using inverse T1w-to-MNI nonlinear transformation. For each participant, an ROI for the parahippocampal place area (PPA) was created by first using an automated metanalyses in Neurosynth with the key term ‘place’. Then, clusters were generated using voxels with a z-score > 2 based on the Neurosynth associative tests. Because these clusters were created through an automated meta-analysis and were not anatomically exclusive to PPA, we visually inspected the results and manually chose the two largest clusters that were spatially consistent with PPA. One cluster was in the left hemisphere (voxel size = 163), and the other cluster was in the right hemisphere (voxel size = 247). These clusters were combined into a single PPA mask. This mask was then transformed into each participant’s native space using the inverse T1w-to-MNI transformation. To create hippocampal ROIs, we used the Automatic Segmentation of Hippocampal Subfields (ASHS)^82^ toolbox with the upenn2020 (10.1016/j.media.2022.102683) atlas to create subfield ROIs in each participant’s hippocampus, including CA3/dentate gyrus (which consisted of CA2, CA3, and dentate gyrus) and CA1. Each participant’s subfield segmentations were manually inspected to ensure the accuracy of the segmentation protocol. These ROIs were then manually edited, for each participant, to only include the hippocampal body. This was done by manually identifying the most anterior and posterior slices of the hippocampal body based on each participant’s T2-weighted anatomical scan. Next, each subfield ROI was transformed into each participant’s native space using the T2-to-T1w transformation, calculated with FLIRT (FSL) with six degrees of freedom, implemented with Nipype. All ROIs were again visually inspected after the transformation to native space to ensure the ROIs were anatomically accurate.

### fMRI pattern similarity scores

fMRI pattern similarity was defined as the Pearson correlation between t-stats maps from a given ROI. All correlations were restricted to t-maps from different scan runs; that is, correlations were never computed between t-maps (trials) from the same scan run. Correlations were performed for all pairwise comparisons of runs and then averaged. For example, to compute the pattern similarity between beach 4 and beach 2 within a given ROI, 56 Pearson correlations were computed, reflecting all possible combinations across different scan runs (e.g., beach 4–run 1 vs. beach 2–run 2, beach 4–run 3 vs. beach 2–run 7, etc.). The 56 correlation coefficients were then Fisher z-transformed and averaged to yield a single similarity value for that scene pair.

For each ROI, pattern similarity between a pair of scenes from the same category (within-category similarity, e.g., beach 4 vs. beach 2) was always expressed relative to the mean pattern similarity between all scenes from opposite categories (across-category similarity). The difference between these measures (within-category minus across-category) is referred to as the *similarity score.* Thus, if a given pair of images from the same category (or a set of images from the same category) has a positive similarity score, this indicates that the brain region (ROI) represented these scenes as more similar than scenes from opposite categories^17,27^. The rationale for using across-category similarity as a baseline is that it facilitates comparisons of representational structure across ROIs. Namely, raw correlations values are difficult to interpret and vary substantially across ROIs. In contrast, *relative* similarity (within- *versus* across-category) corrects for these differences.

### Statistical analyses and reproducibility

#### Split-half reliability for human-based similarity metrics

For human-based measures of memory confusability and perceptual similarity, we conducted reliability analyses. Specifically, split-half reliability was computed separately for each category (beaches, gazebos) as a function of the number of participants included (participant count). For each participant count, 50 random permutations were generated by selecting that number of participants and randomly dividing them into two equal groups. For each participant group, confusability/similarity matrices were averaged across participants. Reliability was defined as the Pearson’s correlation (r) between the matrices for each group. To compute the overall reliability across all 50 permutations, correlation values were first converted to Fisher’s z, then averaged, and then converted back to r (for interpretability).

#### Mixed-effects models

Mixed-effects (LME) models were used to evaluate the extent to which different similarity dimensions, training level, and the interaction between training level and each similarity dimension predicted behavior and/or fMRI similarity scores. The fixed effects (predictors) included in each model are mentioned in the sections below. Critically, in every model that involved similarity dimensions, random intercepts and slopes for similarity dimensions were included for each participant to account for individual differences. Interactions between the identity of the high training category (beaches vs. gazebos) and other fixed effects were included only when they significantly improved model fit or revealed interpretable effects, to avoid overfitting and obscuring lower-order effects. All models were implemented using fitlme or fitglme in MATLAB R2023b, with model comparisons guided by likelihood ratio tests and inspection of residuals to ensure model assumptions were met. Degrees of freedom were computed using the Satterthwaite approximation.

PC scores, memory confusability, perceptual similarity, and normative low training fMRI similarity scores were treated as continuous measures in mixed-effects models. For visualization purposes only, data were also binned (see **Fig 2c**, **Fig. 4c-d**, **Supplementary Fig. 9-11**). For the linear mixed-effects models corresponding to **Fig. 4c,d** and **Supplementary Fig. 11**, the models included both PC1 and PC2 similarity as fixed effects. The lines of fit shown in these figures therefore reflect the effect of one PC while controlling for the other PC. To ensure that the binned visualizations accurately reflect the model, we regressed out PC1 from PC2 (and vice versa) within each scene category before binning the PC scores. In other words, the binned data reflect the effect of one PC while controlling for the other PC. Again, this step (regressing one PC out from the other) was purely for the sake of accurate visualization for the binned data. The same procedure was applied to memory confusability and perceptual similarity in **Supplementary Fig. 9** (i.e., memory confusability was regressed out from perceptual similarity, and vice versa).

#### Establishing behaviorally relevant dimensions of similarity

To identify stimulus similarity dimensions that predicted memory interference errors (**Fig. 2c**), we tested whether PC scores (derived from the 10 metrics of similarity) predicted pairwise confusability in the post-test of the fMRI experiment. Specifically, we ran mixed-effects models for which pairwise confusability was the dependent measure and scores on one of the 10 PCs (**Fig. 1g**) were included as a predictor (separate models for each PC). For each participant, the model included 552 rows corresponding to the 552 pairs of scene images (276 beach pairs and 276 gazebo pairs). If a pair of scenes was confused on the post-test, pairwise confusability for that row was coded as ‘1’; otherwise, confusability was ‘0’ (see **Fig. 2b**). Additional fixed effects included training level (high vs. low), and the designated high-training category for each participant (beaches vs. gazebos).

As an alternate approach, we also directly correlated (Pearson correlation) each subject’s memory confusability matrix, separately for each training condition, with the similarity matrix for each PC. Correlation coefficients were z-transformed, averaged across training conditions within subject, and compared to a test value of 0 using one-sample, two-tailed t-tests.

#### Experience-dependent changes in behavior

To validate the effectiveness of the training manipulation in the fMRI experiment, we analyzed behavioral performance across the training, exposure, and post-test phases using mixed-effects models (**Fig. 3a-c**). Note: these models were based on mean responses per participant/condition (i.e., they did not include trial-level data). For the training phase, two separate models were run. For the first model (**Fig. 3a**, left), the dependent measure was accuracy (% correct) in the test rounds. Fixed effects included training level (high vs. low), training round (round 1 to round 3), and the participant’s designated high-training category (beaches vs. gazebos). This model was used to test for main effects of training round and training level. The second model (**Fig. 3a**, right) focused on errors. For this model, the dependent measure was the frequency of responses and fixed effects included error type (competitor vs. non-competitor), training level (high vs. low), training round (round 1 to round 3), and the participant’s designated high-training category (beaches vs. gazebos). This model was used to test whether interference errors (selecting the competitor) were more common than non-interference errors (selecting the non-competitor). For the exposure phase (**Fig. 3b**) and post-test phase (**Fig. 3c**), the dependent measures in the mixed-effects models were d-prime and accuracy (% correct), respectively and fixed effects included training level (high vs. low) and the participant’s assigned high-training category (beaches vs. gazebos).

To measure the influence of different dimensions of similarity (PC1 vs. PC2) on interference errors during the training phases of the fMRI experiment and the behavioral memory experiment (**Fig. 3d-e**), we used logistic mixed-effects models that included binary trial-by-trial measures of performance. Specifically, the dependent variable in the models represented whether, on each trial, participants made an interference error (selecting the competitor, coded as ‘1’) or correctly selected the target object (coded as ‘0’) (**Fig. 2a, 3a**, right). While rare, trials on which participants selected the non-competitor were excluded from the model. The key fixed effects were the degree centrality of the presented scene (the probe), separately for PC1 and PC2. The rationale for these models was that, in general, higher degree centrality (greater aggregate similarity to other scenes from the same category) would be associated with a higher rate of interference errors. For the fMRI experiment (**Fig. 3d**), other fixed effects included in the models were the training level (high vs. low), training round (round 1 to round 3), and the designated high-training category for the participant (beaches vs. gazebos). For the behavioral memory experiment (**Fig. 3e**), other factors included in the models were the training round (round 1 to round 3), and the category of the current probe (beach vs. gazebo).

To formally compare interaction effects with training across PC1 and PC2, we performed a linear contrast test (F-test) on the associated fixed effects of our model to evaluate the significance of coefficient differences (i.e., H_0_: β_(PC1 × training)_ − β_(PC2 × training)_ = 0 for the fMRI experiment, H_0_: β_(PC1 × round)_ − β_(PC2 × round)_ = 0 for the behavioral memory experiment).

#### Relationships between experience, similarity dimensions, and the hippocampus

To test the overall effects of training on fMRI similarity scores for each ROI (**Fig. 4b**), we used linear mixed-effects models where the dependent measure was the mean fMRI similarity score, and fixed effects included training level (high vs. low) and the participant’s designated high-training category (beaches vs. gazebos). Note: these models were based on mean responses per participant/condition (i.e., they did not include trial-level data). In addition, to test whether an ROI’s average similarity score was above zero, two-tailed one-sample t-tests (contrast value = zero) were conducted.

To test whether fMRI similarity scores within each ROI were related to dimensions of visual similarity (PC1, PC2 for **Fig. 4c-d** and **Supplementary Fig. 11**; all 10 PCs for **Supplementary Table 3**) and training level (low vs. high), we ran a separate set of linear mixed-effects models. In these models, the dependent measure was the fMRI similarity score for every pairwise combination of scenes from the same category (baselined against the mean across-category similarity). Specifically, for each participant, the model included 552 fMRI similarity values, reflecting every pairwise combination of beaches (276 total) and gazebos (276 total). Fixed effects included PC scores (continuous predictors), training level (low vs. high), and the high-training scene category for each participant (beaches vs. gazebos).

To further support the argument that repulsion within CA3/DG was a direct reaction to memory interference, we implemented another mixed-effects model that predicted similarity scores in CA3/DG, but instead of using PC1 and PC2 scores as fixed effects, we used similarity values taken directly from the two behavioral similarity experiments (**Fig. 1c**): the behavioral memory experiment and the perceptual similarity experiment (**Supplementary Fig. 9**). In these models, the dependent measure was CA3/DG similarity scores (552 values per participant) and fixed effects included perceptual similarity and memory confusability values (continuous predictors), training level (low vs. high), and the high-training scene category for each participant (beaches vs. gazebos).

Finally, for each ROI and each scene category, we tested the extent to which the representational structure after high training preserved vs. transformed the representational structure after low training (independent of any similarity metric or PC scores) (**Supplementary Fig. 10**). Because the assignment of each scene category to each training condition was counterbalanced *across participants*, this analysis required across-participant alignment. Specifically, for each participant, pairwise similarity scores in the high training condition were treated as the dependent measure whereas similarity scores in the low training condition *from other participants* were the critical predictor (fixed effect). For example, for a given participant, if the high training category was beaches, then the dependent measure for that participant would be the 276 pairwise similarity scores for beaches. To obtain the corresponding similarity scores for the low training condition, the mean similarity score (for each of the 276 pairs of beaches) was computed from all participants (N = 26) for which the low training category was beaches. Thus, participant-specific and category-specific representational structure in the high training condition was considered as a function of *normative representational structure* in the low training condition. We refer to this normative measure as the ‘normative low-training fMRI similarity scores’. For the mixed-effects models, the high-training category (beaches vs gazebos) was also included as a fixed effect.

#### Statistical significance

A threshold of p < .05 was used to determine statistical significance. All t-tests were two-tailed. To correct for multiple comparisons when testing effects across all 10 PCs (**Fig. 2c**, **Supplementary Table 3**), we applied a Bonferroni-adjusted significance threshold of α = 0.005.

## Data availability

All data and materials will be available by the time of publication at https://osf.io/gy23h/. Source Data are provided with this paper.

## Code availability

Analysis codes used for the analysis will be available by the time of publication at https://osf.io/gy23h/.

## Acknowledgements

This work was supported by National Institutes of Health grant NIH-NINDS 2R01NS089729 to B.A.K and Ruth L. Kirschstein NRSA Individual Predoctoral Fellowship (from the National Institutes of Health) grant F31NS126016 to W.G.

## Author contributions

Conceptualization: S.M., W.G., D.G., U.M., B.A.K. Methodology: S.M., W.G., D.G., U.M., E.W., B.A.K. Investigation: S.M., W.G., D.G., B.A.K. Visualization: S.M., B.A.K. Funding acquisition: W.G., B.A.K. Project administration: U.M., B.A.K. Supervision: U.M., B.A.K. Writing, first draft: S.M., B.A.K. Review and editing: S.M., W.G., D.G., B.A.K.

## Additional information

**Correspondence and requests for materials** should be addressed to Soroush Mirjalili or Brice A. Kuhl

---

**Supplementary Fig. 1.**
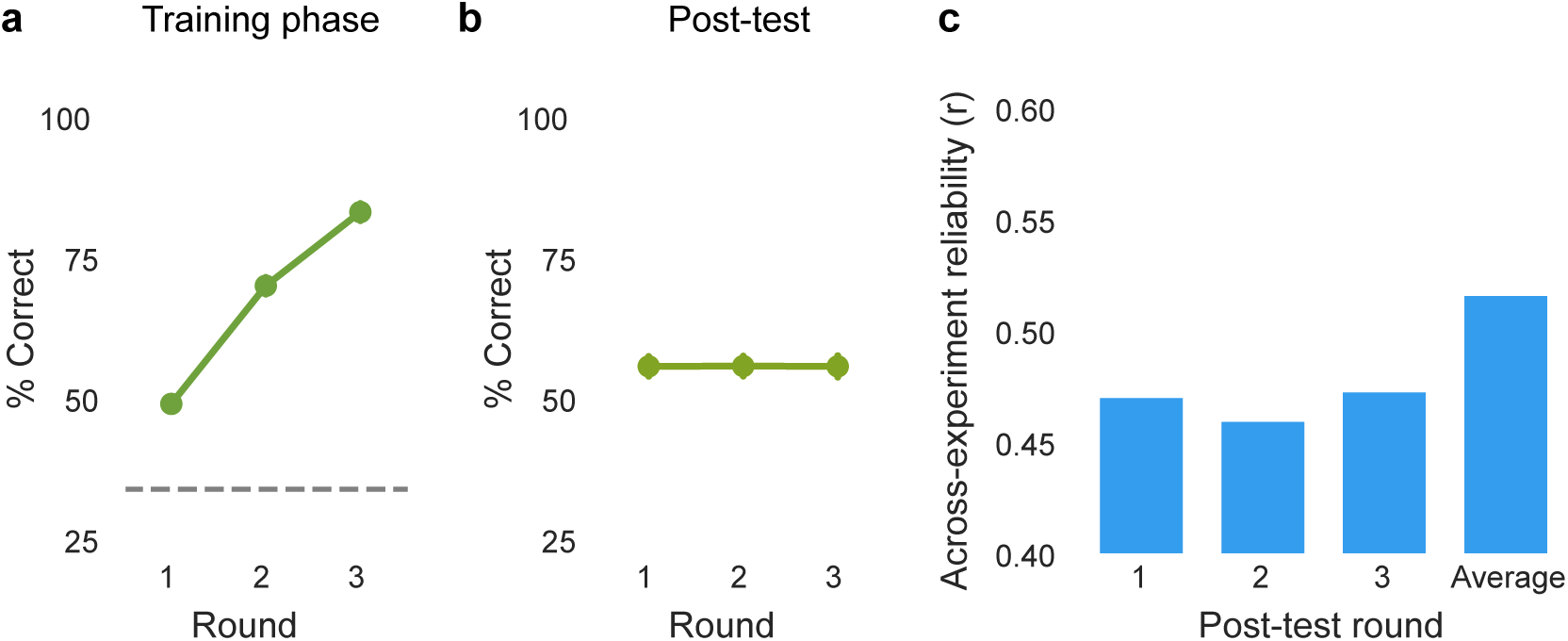
Performance in the behavioral memory experiment. **a**, Performance (percent current) during each test round of the training phase. **b**, Performance (percent correct) for each round of the post-test. Note: no additional study trials or feedback occurred between the three post-test rounds. **c**, Pearson correlations between memory confusability in the fMRI experiment and memory confusability in the behavioral memory experiment. For each experiment, a confusion matrix was derived from the post-test, averaging across all participants. For the behavioral memory experiment, we derived the confusion matrix using only post-test data from round 1, only from round 2, only from round 3, or using data averaged across all three rounds. Similarity between the experiments was highest when the post-test data from the behavioral experiment were averaged across rounds.

**Supplementary Fig. 2.**
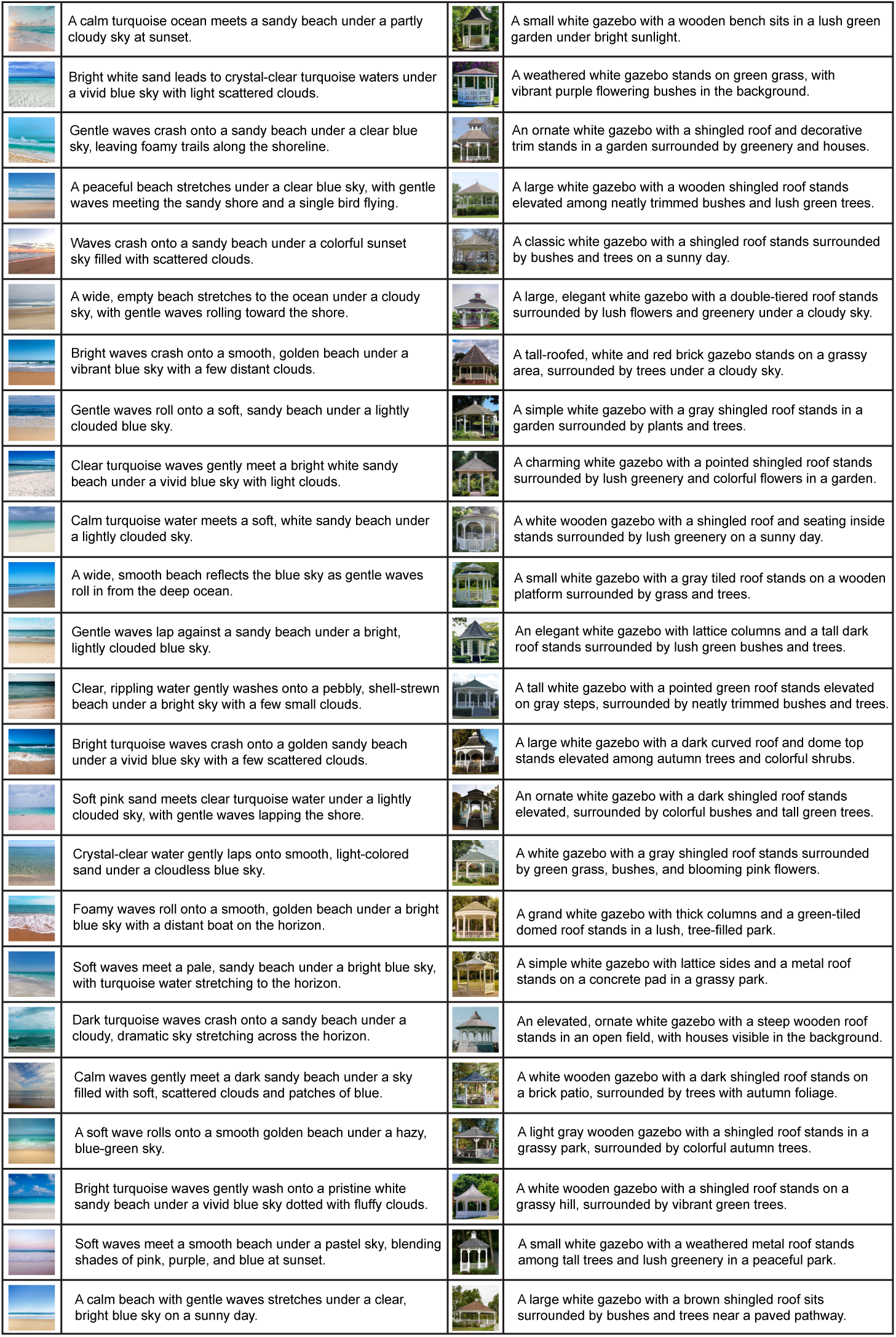
Text descriptions, generated by OpenAI’s ChatGPT (GPT-4-turbo; gpt-4o variant), for each scene image. Text descriptions were used for the Natural Language Processing (NLP) of text similarity metric. We performed a follow-up analysis to assess the extent to which NLP-based text similarity was driven by nouns versus adjectives in the scene descriptions. Specifically, we computed text-similarity matrices using three versions of the descriptions: the original descriptions, nouns only, and adjectives only. Pairwise Pearson correlations between the resulting similarity matrices were as follows: original vs. nouns only, r = 0.426; original vs. adjectives only, r = 0.299; nouns only vs. adjectives only, r = 0.237.

**Supplementary Fig. 3.**
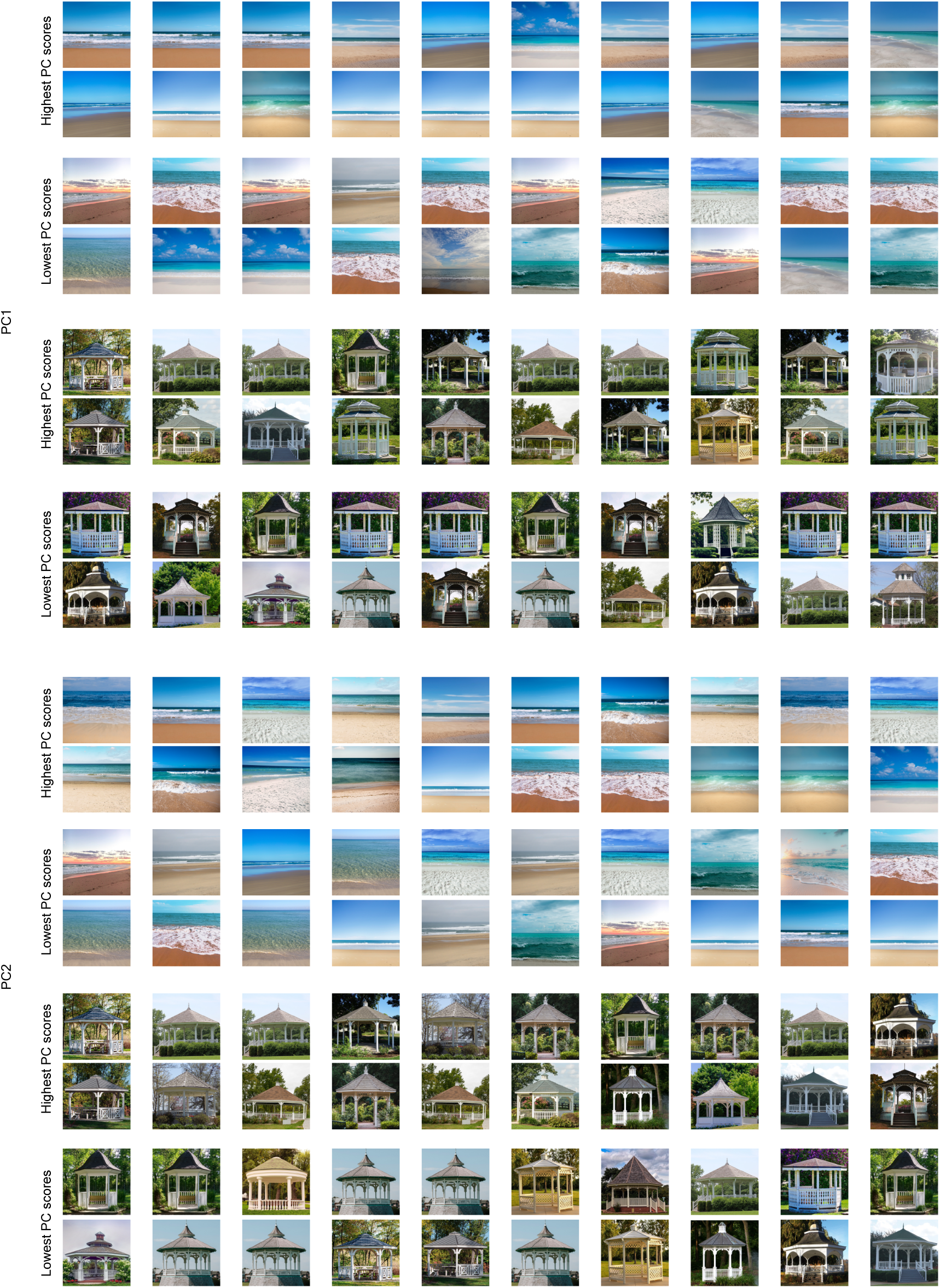
Scene pairs with highest/lowest PC scores (PC1 and PC2). Pairs are arranged vertically, separately for each PC and scene category. Pairs with the 10 highest/lowest scores are shown (from left to right: descending order for highest, ascending order for lowest).

**Supplementary Fig. 4.**
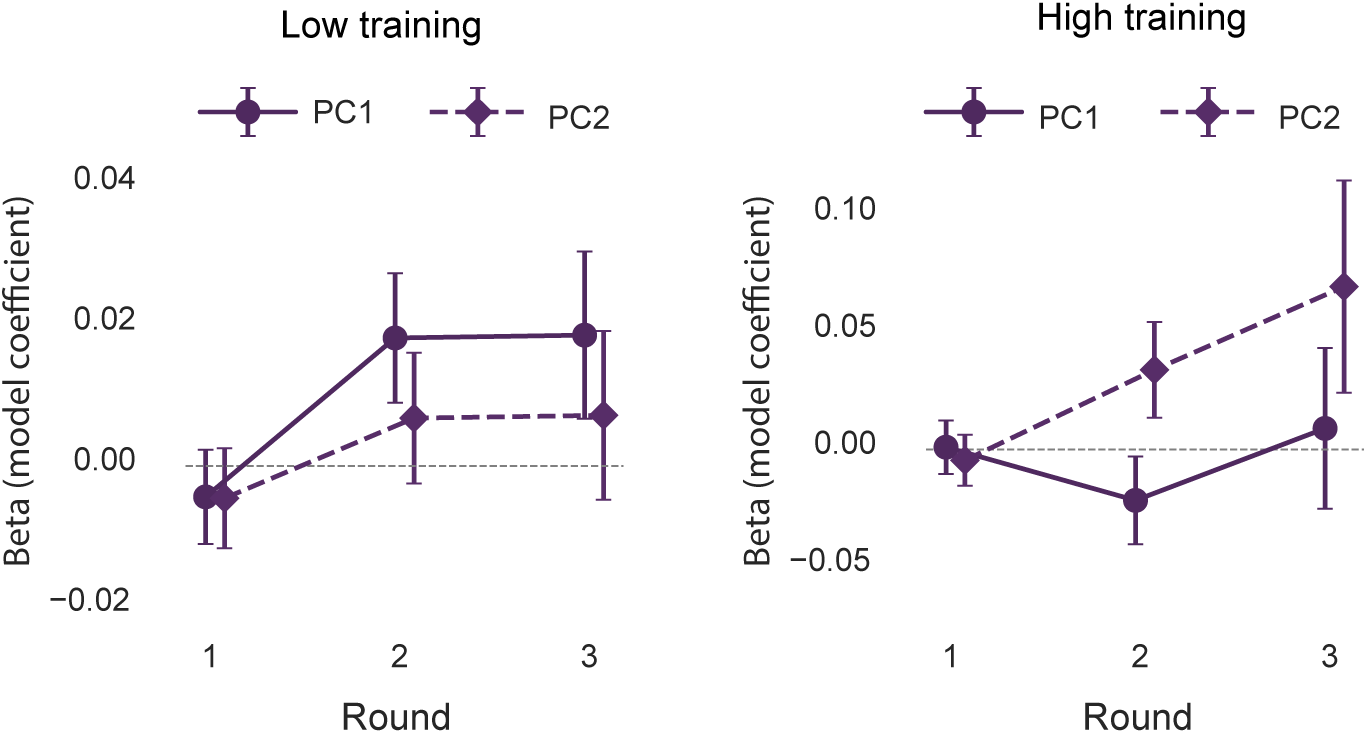
The influence of each similarity dimension (PC1, PC2) on interference errors during each round of the training phase of the fMRI experiment. For the low training condition (Left) and the high training condition (Right), we used mixed-effects models in which the dependent variable represented trial-by-trial interference errors (selecting the competitor; Fig. 2a). The key independent fixed effects were the degree centrality of the presented scene, separately for PC1 and PC2, and round of training. With low training, for PC1, there was a significant interaction between degree centrality and round of training (*β* = 0.0137, t_(3732)_ = 2.491, P = 0.013), while there was no effect for PC2 (*β* = 0.0019, t_(3732)_ = 0.338, P = 0.735). In contrast, with high training, for PC2, there was a significant interaction between degree centrality and round of training (*β* = 0.0392, t_(3732)_ = 2.528, P = 0.012), while there was no effect for PC1 (*β* = −0.0039, t_(3732)_ = −0.272, P = 0.786). Circle (PC1) and diamond (PC2) markers indicate beta estimates from linear mixed-effects models in which trial-by-trial degree centrality values (from PC1 and PC2) were used to predict the likelihood of an interference error (selecting the competitor). Error bars represent the corresponding S.E.M.s from the models.

**Supplementary Fig. 5.**
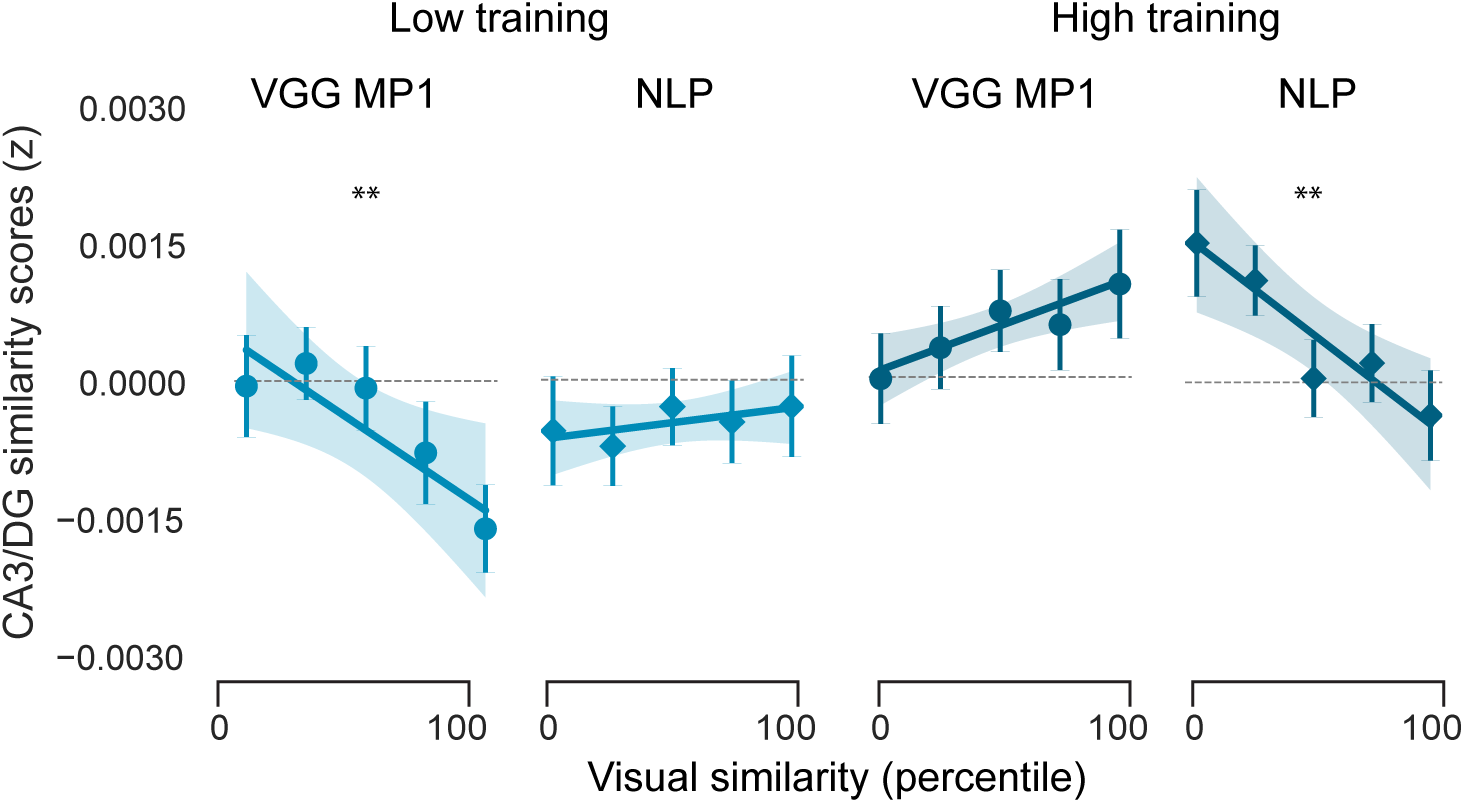
Effects of VGG MP1 and NLP of text similarity on CA3/DG similarity scores. To identify the metrics of similarity that best captured perceptual vs semantic similarity, we asked ChatGPT the following: ‘We have 8 measures of similarity, driven by various computational models. The measures include 1-CLIP similarity which is image-based completely, 2- NLP of text descriptions of the scenes, 3-VGG MP1, 4-VGG MP3, 5-VGG MP5, 6-AlexNet, last fully connected layer, 7- GIST, 8-SSIM. Which one should I use for a perceptual similarity measure and which for a semantic similarity measure?’ ChatGPT’s response was: ‘My recommendation: Best perceptual model: Use VGG pool1. VGG pool1 is often the most defensible neuroscience choice. Best semantic model: use annotation embeddings if you want the cleanest conceptual semantics independent of vision.’ We next ran a linear mixed-effects model with pairwise CA3/DG similarity scores (552 pairs per participant) as the dependent variable. The predictors (fixed effects) included VGG MP1 similarity (as the perceptual similarity), NLP of text similarity (as the semantic similarity), training (low vs. high), and the identity of the high-training category (beaches vs. gazebos). For, each similarity metric, there was a highly significant interaction between stimulus similarity and training level (VGG MP1: *β* = 0.0109, t_(28697)_ = 3.520, P < 0.001; NLP of text: *β* = −0.0130, t_(28697)_ = −4.657, P < 0.001). However, these interactions took opposite forms. With low training, CA3/DG similarity was negatively related to VGG MP1 similarity (*β* = −0.0075, t_(14348)_ = −3.166, P = 0.002), but was not influenced by NLP of text similarity (*β* = 0.0033, t_(14348)_ = 0.475, P = 0.475). With high training, however, CA3/DG similarity was not influenced by VGG MP1 similarity (*β* = 0.0036, t_(14348)_ = 1.407, P = 0.160), but was negatively related to NLP of text similarity (*β* = −0.0099, t_(14348)_ = -2.647, P = 0.008). In the figure, circles represent VGG MP1 effects and diamonds represent NLP of text effects. For both metrics, similarity was divided into five bins, solely for visualization purposes. Each circle or diamond marker represents the mean similarity score within a given similarity bin, and the accompanying error bar indicates S.E.M. Lines of best fit—established from linear mixed-effects models—are shown for reference, with shaded areas representing the S.E.M (**P < 0.01).

**Supplementary Fig. 6.**
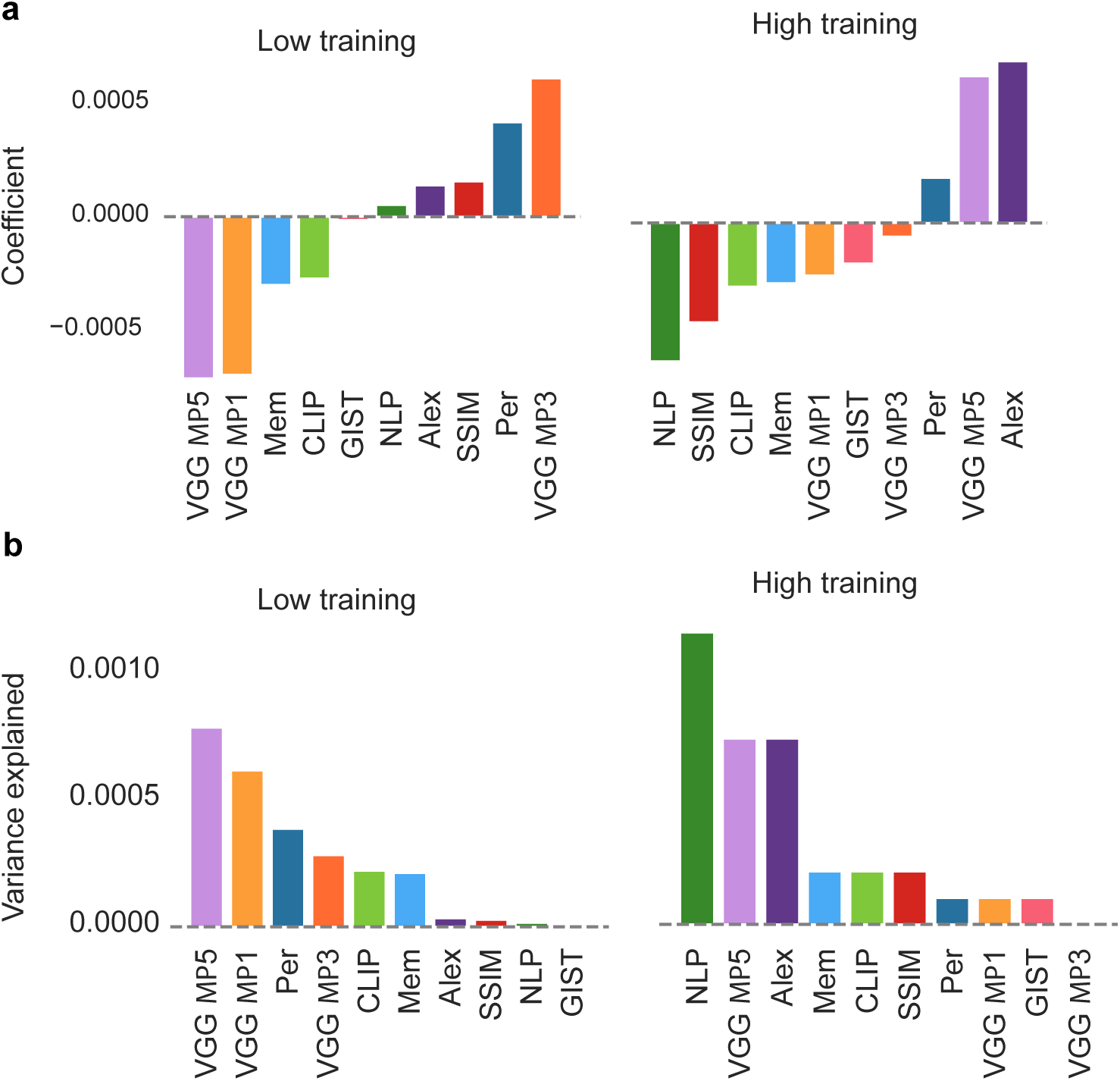
Influence of each metric of similarity on CA3/DG similarity. **a**, Regularized regression using ridge (L2) penalization was performed in which the 10 metrics of similarity were used to predict CA3/DG similarity, separately for low and high training. This approach allowed us to estimate the relative contribution of each similarity metric to CA3/DG similarity while stabilizing coefficient estimates. The model coefficients are shown for the low training condition (left), and high training condition (right), rank ordered by coefficient value. **b**, Variance partitioning was performed to characterize the relative contribution of each of the 10 similarity metrics to CA3/DG similarity. Specifically, we ran linear mixed-effects models, separately for low and high training, with pairwise CA3/DG similarity scores as the dependent variable. The predictors (fixed effects) included all 10 metrics of similarity. We then compared the variance explained by the full model to that of reduced models in which each predictor was omitted in turn. Plots show the unique variance explained by each similarity metric (rank ordered from most to least), separately for the low training condition (left) and high training condition (right).

**Supplementary Fig. 7.**
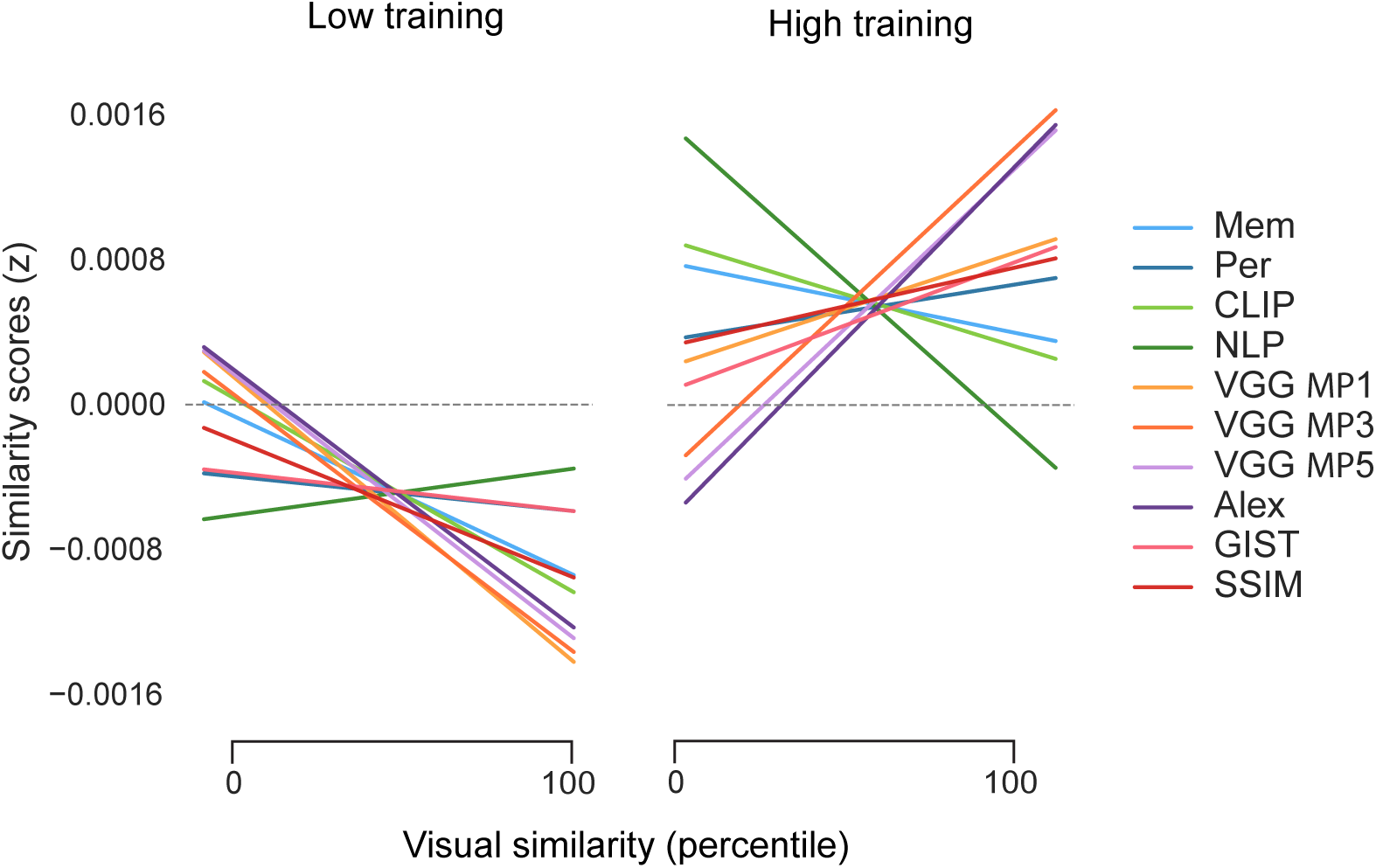
Relationships between each similarity metric and CA3/DG similarity. For each of the 10 metrics of similarity, and separately for the low training condition (left) and high training condition (right), we ran a linear mixed-effects model (20 models in total) with pairwise CA3/DG similarity scores (276 pairs per training condition per participant) as the dependent variable. The predictors (fixed effects) included only one of the 10 metrics of similarity and the identity of the high-training category (beaches vs. gazebos). Lines of best fit—established from linear mixed-effects models—are shown for reference.

**Supplementary Fig. 8.**
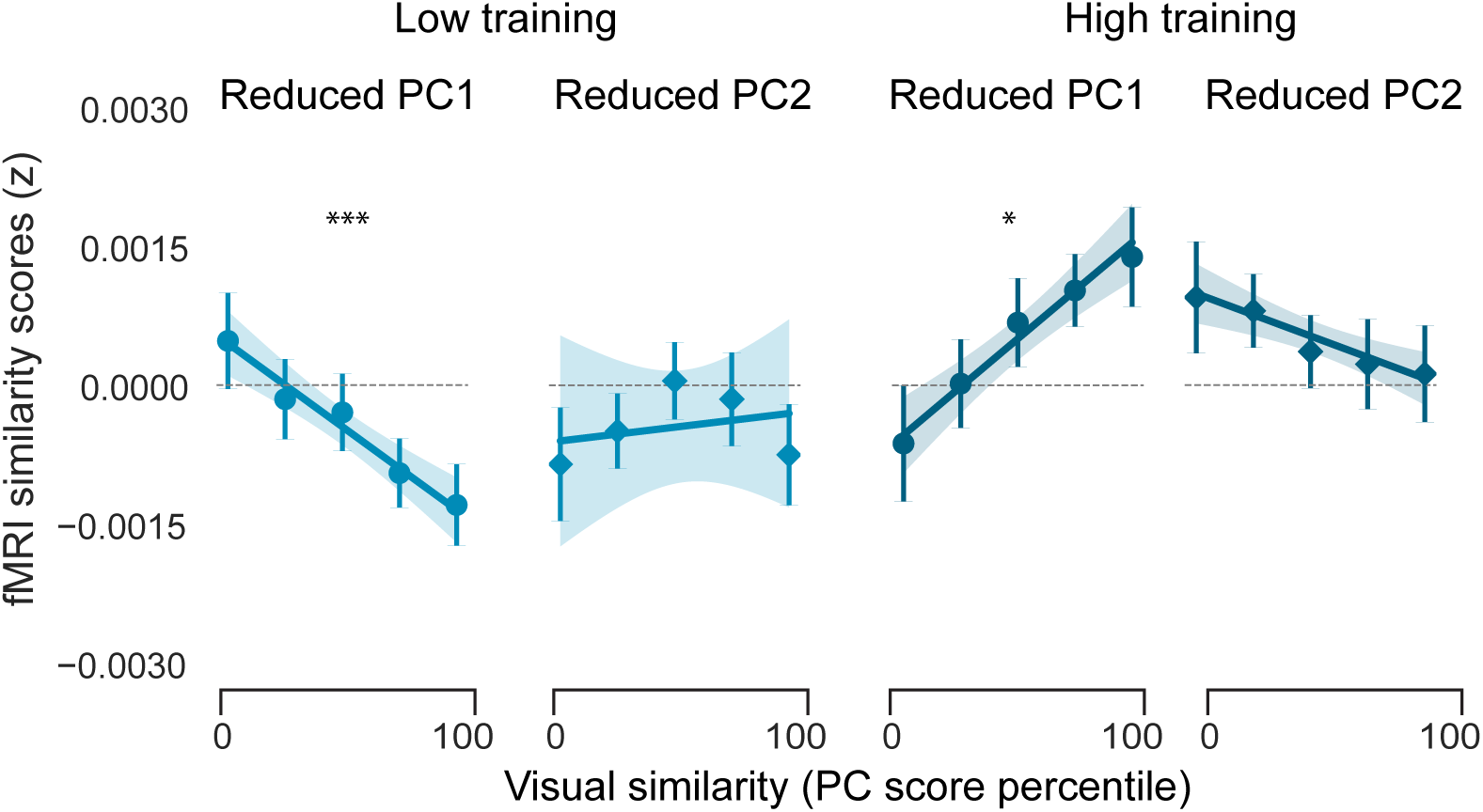
Relationships between PC scores and CA3/DG similarity after removing similarity metrics based on human behavior. After excluding the two human-based similarity metrics (memory confusability and perceptual similarity), the PCA was recomputed, such that the resulting dimensions were based entirely on computational (non-human) metrics. We then repeated the analyses shown in Fig. 4c using the PC1 and PC2 scores obtained from this reduced PCA model. Specifically, we ran a linear mixed-effects model with pairwise CA3/DG similarity scores (552 pairs per participant) as the dependent variable. The predictors (fixed effects) included the reduced PC1 and PC2 scores, training (low vs. high), and the identity of the high-training category (beaches vs. gazebos). The interaction between PC score and training level was highly significant for both PC1 and PC2 (PC1: *β* = 0.0018, t_(28697)_ = 7.580, P < 0.001; PC2: *β* = −0.0006, t_(28697)_ = -2.688, P = 0.007). However, these interactions took opposite forms. With low training, CA3/DG similarity was negatively related to PC1 similarity (*β* = −0.0009, t_(14348)_ = −4.278, P < 0.001), but there was no effect of PC2 similarity (*β* = 0.0003, t_(14348)_ = 0.751, P = 0.453). With high training, CA3/DG similarity was positively related to PC1 similarity (*β* = 0.0010, t_(14348)_ = 2.529, P = 0.011), but there was no effect of PC2 similarity (*β* = - 0.0004, t_(14348)_ = −1.105, P = 0.269). In the figure, circles represent reduced PC1 effects and diamonds represent reduced PC2 effects. For both PCs, similarity was divided into five bins, solely for visualization purposes. Each circle or diamond marker represents the mean similarity score within a given similarity bin, and the accompanying error bar indicates S.E.M. Lines of best fit—established from linear mixed-effects models—are shown for reference, with shaded areas representing the S.E.M (*P < 0.05, ***P < 0.001).

**Supplementary Fig. 9.**
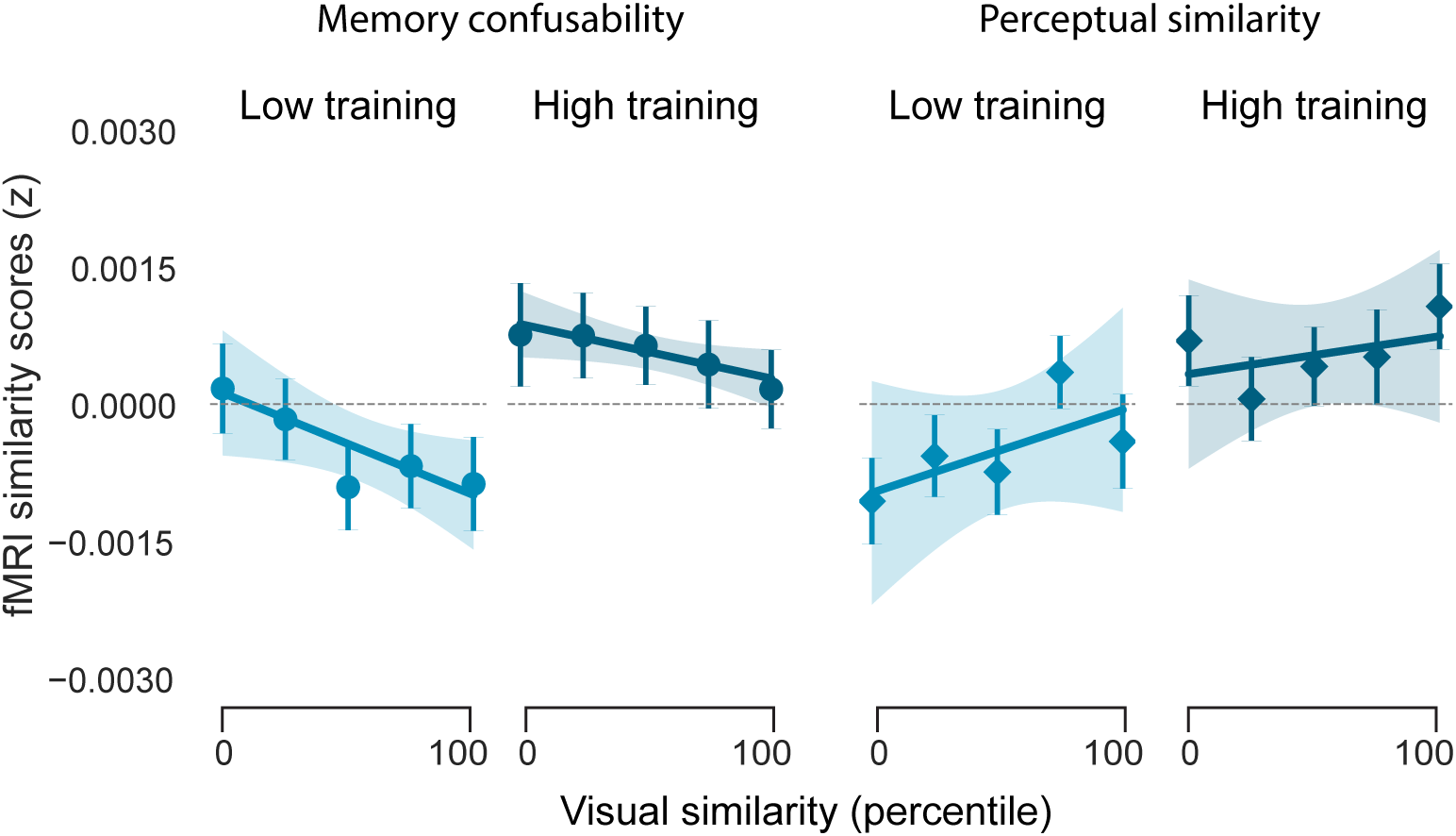
Effects of memory confusability and perceptual similarity on CA3/DG similarity scores. For the two human-based measures of similarity (memory confusability and perceptual similarity judgments), we tested for relationships with similarity scores in CA3/DG. Given the relevance of CA3/DG to memory, we hypothesized that CA3/DG similarity scores would specifically be sensitive to memory confusability. We ran a linear mixed-effects model with pairwise CA3/DG similarity scores (552 pairs per participant) as the dependent variable. The predictors (fixed effects) included memory confusability, perceptual similarity, their interaction with training (low vs. high), and the identity of the high-training category (beaches vs. gazebos). There was a significant negative main effect of memory confusability (β = −0.00062, t_(28692)_= -2.157, P = 0.031), with a qualitatively positive main effect of perceptual similarity (β = 0.00069, t_(28692)_ = 1.390, P = 0.165). There was no interaction with training with either measure of similarity (β’s < 0.00016, t_(28692)_ ’s < 0.427, Ps > 0.669). Thus, CA3/DG inverted similarity as defined by memory confusability, but not as defined by perceptual judgments. The fact that memory confusability did not interact with training level (which contrasts with effects for PC1 and PC2 shown in Fig. 4c) is presumably because the memory confusability metric captured dimensions of similarity that were relevant across training levels. Indeed, memory confusability positively loaded on both PC1 and PC2 (see Fig. 1i and **Supplementary Table 1**). Note: the order of the plots differs from Fig. 4c to better illustrate the consistency of effects across training conditions. In the figure, circles represent memory confusability effects and diamonds represent perceptual similarity effects. For both metrics, similarity was divided into five bins, solely for visualization purposes. Each circle or diamond marker represents the mean similarity score within a given similarity bin, and the accompanying error bar indicates S.E.M. Lines of best fit—established from linear mixed-effects models—are shown for reference, with shaded areas representing the S.E.M. Additionally, we performed a formal model comparison analysis. Specifically, we compared a baseline mixed-effects model predicting CA3/DG similarity from perceptual similarity alone and training against an expanded model that additionally included memory confusability. A likelihood ratio test revealed that inclusion of memory confusability significantly improved model fit (χ²_(1)_ = 8.860, P = 0.003). These results indicate that memory confusability explains unique variance in hippocampal representations beyond perceptual similarity alone.

**Supplementary Fig. 10.**
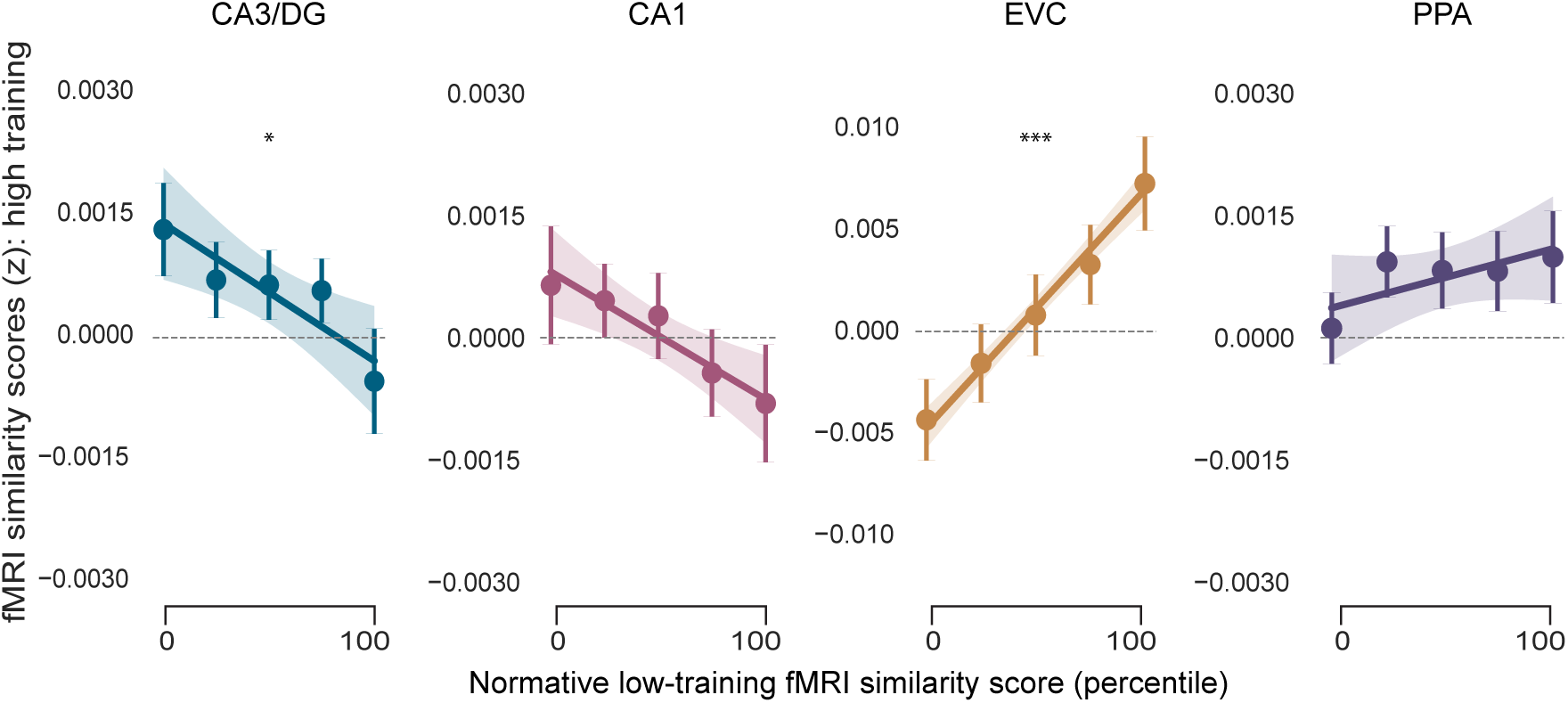
Transformation of representational structure across training levels. Here, we tested the extent to which the representational structure associated with high training preserved vs. transformed the representational structure associated with low training, independent of any similarity metric or dimension (see Methods – ‘Relationships between experience, similarity dimensions, and the hippocampus’). For each ROI, we ran linear mixed-effects models for which the dependent measure was each participant’s fMRI similarity scores for the high-training category (276 pairwise values). The key predictor (fixed effect) was normative low-training fMRI similarity scores. The normative low-training fMRI similarity scores reflected the mean similarity score for each scene pair across all *other* participants for which that scene category was associated with low training. The identity of the high-training category (beaches vs gazebos) was included as a fixed effect. The relationship between normative low-training similarity scores and (participant specific) high-training similarity scores was negative for CA3/DG (β = −0.184, t_(14348)_= −1.970, P = 0.049), but strongly positive for EVC (β = 0.756, t_(14348)_= 10.408, P < 0.001). Effects were not significant for CA1 (β = −0.123, t_(14348)_= −1.468, P = 0.142) or PPA (β = 0.072, t_(14348)_= 0.899, P = 0.369). Thus, whereas EVC robustly *preserved* within-category representational structure across stages of learning, CA3/DG *transformed* representational structure across stages of learning. In the figure, normative low-training fMRI similarity scores are divided into five bins, solely for visualization purposes. Each circle marker represents the mean similarity score within a given similarity bin, and the accompanying error bar indicates S.E.M.. Lines of best fit—established from linear mixed-effects models—are shown for reference, with shaded areas representing the S.E.M (*P < 0.05, ***P < 0.001).

**Supplementary Fig. 11.**
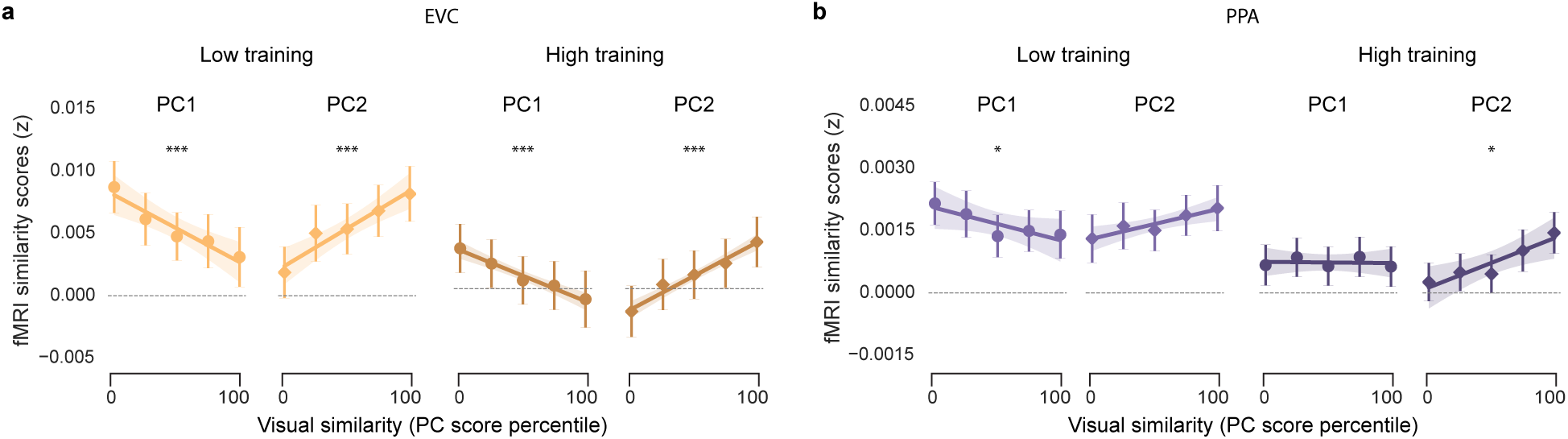
Relationships between visual similarity, training level, and fMRI similarity scores in visual cortical regions. We tested whether fMRI similarity scores in early visual cortex (EVC, **a**) and parahippocampal place area (PPA, **b**) were related to visual similarity on each dimension (PC1 and PC2 scores) and/or training level (low vs. high training). This was tested via linear mixed-effects models that were identical, in structure, to the models used for CA3/DG and CA1 (Fig. 4c**-d**). **a,** For EVC, there were significant main effects of PC1 (β = −0.0018, t_(28692)_ = −4.812, P < 0.001) and PC2 (β = 0.0022, t_(28692)_ = 6.461, P < 0.001), however these effects were in opposite directions (negative effect for PC1, positive effect for PC2). While the effect for PC1 was qualitatively similar across training levels (negative in both cases: β’s < −0.0014, t_(14348)_’s < −3.619, P’s < 0.001), there was a significant interaction between training level and PC1 (β = 0.00070, t_(28692)_ = 3.437, P = 0.005). The effect for PC2 was positive across training levels (β’s > 0.0020, t_(14348)_’s > 5.777, P’s < 0.001) and there was no interaction between training level and PC2 (β = −0.00038, t_(28692)_ = −1.914, P = 0.056). **b**, For PPA, there was a significant main effect for PC2 (β = 0.00037, t_(28692)_ = 2.589, P = 0.010), but not PC1 (β = −0.00016, t_(28692)_ = −1.158, P = 0.247). There was a significant interaction between training and PC1 (β = 0.00029, t_(28692)_ = 2.636, P = 0.008) but no significant interaction between training and PC2 (β = 0.00018, t_(28692)_ = 1.666, P = 0.096). For PC1, there was a negative effect of PC score for the low training condition (β = −0.00031, t_(14348)_ = -2.491, P = 0.013) and no effect in the high training condition (β = 0.00001, t_(14348)_ = −0.106, P = 0.915). For PC2, there was no effect of PC score in the low training condition (β = 0.00028, t_(14348)_ = 1.677, P = 0.094), but there was a positive effect for the high training condition (β = 0.00046, t_(14348)_ = 2.971, P = 0.003). In **a** and **b**, circles represent effects of PC1 and diamonds represent effects of PC2. For visualization purposes, PC scores are sorted and divided into five bins, as in Fig. 2c. Each circle or diamond marker represents the mean similarity score within a given similarity bin, and the accompanying error bar indicates S.E.M. Lines of best fit—established from linear mixed-effects models—are shown for reference, with shaded areas representing S.E.M. Asterisks above plots reflect the main effect of PC scores obtained from linear mixed-effects models (*P < 0.05, ***P < 0.001).

**Supplementary Fig. 12.**
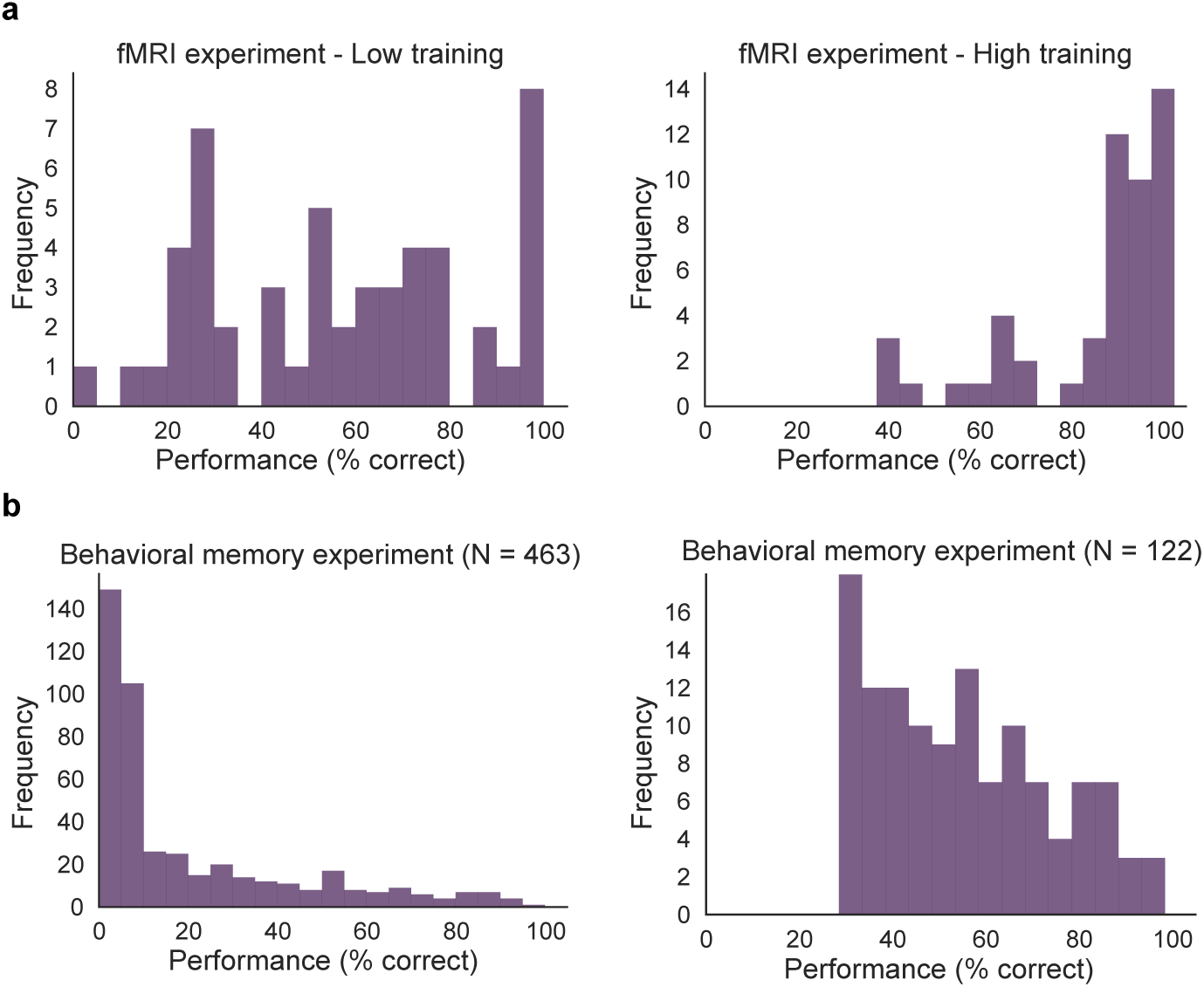
Distribution of post-test performance for the fMRI and behavioral memory experiments. **a**, Histograms of participant-level performance (overall percent correct) for the low training condition (Left) and high training condition (Right). **b**, Histograms of participant-level performance (overall percent correct, averaged over the three rounds of post-test) for the behavioral memory experiment before performance-based exclusions were applied (Left) and after a performance cutoff was applied (Right). The cutoff for the behavioral memory experiment was percent correct > 28%. The rationale for applying a performance cutoff was to increase the reliability of confusion matrices. While the cutoff resulted in the exclusion of more than 73.6% of the participants in the behavioral memory experiment, the excluded participants had lower performance than almost all of the participants in the fMRI experiment.

**Supplementary Table 1.** Loadings for all 10 Principal Components (PCs) from the Principal Component Analysis described in Fig. 1. Rows correspond to the PC number and columns correspond to each of the 10 similarity metrics.

|  | Memory | Percept. | CLIP | NLP | VGG MP1 | VGG MP3 | VGG MP5 | Alex. | GIST | SSIM |
| --- | --- | --- | --- | --- | --- | --- | --- | --- | --- | --- |
| **PC1** | 0.240 | 0.191 | 0.236 | -0.081 | 0.427 | 0.442 | 0.332 | 0.426 | -0.127 | 0.399 |
| **PC2** | 0.438 | 0.515 | 0.272 | 0.482 | -0.106 | -0.146 | 0.113 | 0.005 | 0.325 | -0.293 |
| **PC3** | -0.236 | -0.124 | -0.252 | -0.154 | -0.064 | 0.166 | 0.534 | 0.104 | 0.705 | -0.128 |
| **PC4** | -0.410 | -0.443 | 0.570 | 0.537 | 0.038 | 0.082 | 0.118 | -0.015 | 0.007 | 0.020 |
| **PC5** | -0.017 | 0.044 | 0.689 | -0.662 | -0.157 | -0.119 | -0.064 | -0.005 | 0.139 | -0.150 |
| **PC6** | 0.615 | -0.511 | 0.044 | -0.023 | 0.146 | 0.176 | -0.154 | -0.408 | 0.304 | 0.154 |
| **PC7** | -0.369 | 0.365 | 0.028 | 0.004 | 0.568 | 0.131 | -0.412 | -0.315 | 0.343 | 0.068 |
| **PC8** | 0.093 | -0.278 | -0.062 | 0.022 | 0.277 | -0.049 | -0.425 | 0.665 | 0.148 | -0.432 |
| **PC9** | 0.057 | -0.087 | -0.014 | -0.101 | 0.545 | -0.192 | 0.432 | -0.284 | -0.310 | -0.531 |
| **PC10** | -0.019 | 0.081 | -0.005 | -0.007 | -0.245 | 0.805 | -0.116 | -0.140 | -0.183 | -0.467 |

**Supplementary Table 2.** Robustness of PC1 and PC2 to removal of individual similarity metrics. The original Principal Component Analysis (PCA) that was used to identify PC1 and PC2 in the main text was repeated after removing one similarity metric at a time, resulting in a 552 × 9 input matrix for each reduced model. The table shows the correlation coefficients from Pearson correlations between the original PC1 and PC2 scores and the ‘new’ PC scores for each reduced model across the 552 scene pairs.

| Excluded Metric | New vs original PC1 | New vs original PC2 |
| --- | --- | --- |
| Memory confusability | 0.991 | 0.935 |
| Perceptual similarity | 0.994 | 0.924 |
| CLIP | 0.993 | 0.979 |
| NLP of text | 0.999 | 0.940 |
| VGG MP1 | 0.990 | 0.996 |
| VGG MP3 | 0.987 | 0.990 |
| VGG MP5 | 0.988 | 0.994 |
| AlexNet | 0.991 | 1.000 |
| GIST | 0.998 | 0.960 |
| SSIM | 0.981 | 0.972 |

**Supplementary Table 3.**
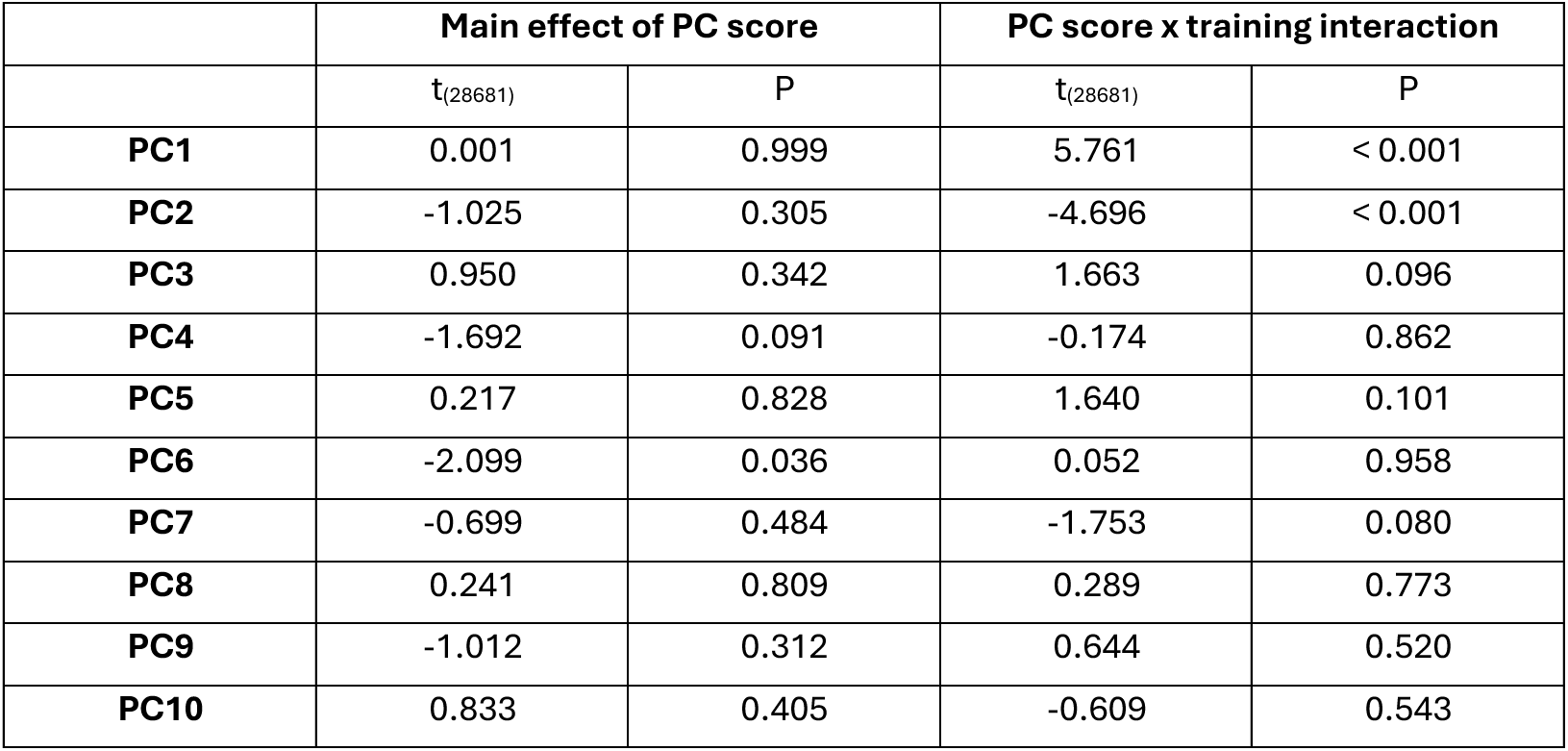
Relationships between visual similarity dimensions, training level, and similarity scores in CA3/DG. Whereas the linear mixed effect model related to Fig. 4c included two dimensions of visual similarity (PC1, PC2) and training level (low vs. high), here we ran an expanded model that included all 10 dimensions of similarity (PC1 – PC10 scores). Each row in the table shows the main effect of PC score (for each similarity dimension) as well as the interaction between PC scores (for each similarity dimension) and training level. As in Fig. 4c, the model included a fixed effect representing the identity of the high-training category (beaches vs. gazebos). To correct for multiple comparisons (separately for main effects and interactions), we used a Bonferroni-adjusted significance threshold of α = 0.005. At this threshold, the only significant effects were interactions between PC score and training level for PC1 and PC2.

|  | Main effect of PC score |  | PC score x training interaction |  |
| --- | --- | --- | --- | --- |
| | $t_{(28681)}$ | P | $t_{(28681)}$ | P |
| **PC1** | 0.001 | 0.999 | 5.761 | < 0.001 |
| **PC2** | -1.025 | 0.305 | -4.696 | < 0.001 |
| **PC3** | 0.950 | 0.342 | 1.663 | 0.096 |
| **PC4** | -1.692 | 0.091 | -0.174 | 0.862 |
| **PC5** | 0.217 | 0.828 | 1.640 | 0.101 |
| **PC6** | -2.099 | 0.036 | 0.052 | 0.958 |
| **PC7** | -0.699 | 0.484 | -1.753 | 0.080 |
| **PC8** | 0.241 | 0.809 | 0.289 | 0.773 |
| **PC9** | -1.012 | 0.312 | 0.644 | 0.520 |
| **PC10** | 0.833 | 0.405 | -0.609 | 0.543 |

